# The transmembrane domain of the desmosomal cadherin desmoglein-1 governs lipid raft association to promote desmosome adhesive strength

**DOI:** 10.1101/2024.04.24.590936

**Authors:** Stephanie E. Zimmer, William Giang, Ilya Levental, Andrew P. Kowalczyk

**Affiliations:** Departments of Dermatology and Cellular and Molecular Physiology, Pennsylvania State University College of Medicine, Hershey, PA, USA; Department of Molecular Physiology and Biological Physics, University of Virginia, Charlottesville, VA, USA

## Abstract

Cholesterol- and sphingolipid-enriched domains called lipid rafts are hypothesized to selectively coordinate protein complex assembly within the plasma membrane to regulate cellular functions. Desmosomes are mechanically resilient adhesive junctions that associate with lipid raft membrane domains, yet the mechanisms directing raft association of the desmosomal proteins, particularly the transmembrane desmosomal cadherins, are poorly understood. We identified the desmoglein-1 (DSG1) transmembrane domain (TMD) as a key determinant of desmoglein lipid raft association and designed a panel of DSG1_TMD_ variants to assess the contribution of TMD physicochemical properties (length, bulkiness, and palmitoylation) to DSG1 lipid raft association. Sucrose gradient fractionations revealed that TMD length and bulkiness, but not palmitoylation, govern DSG1 lipid raft association. Further, DSG1 raft association determines plakoglobin recruitment to raft domains. Super-resolution imaging and functional assays uncovered a strong relationship between the efficiency of DSG1_TMD_ lipid raft association and the formation of morphologically and functionally robust desmosomes. Lipid raft association regulated both desmosome assembly dynamics and DSG1 cell surface stability, indicating that DSG1 lipid raft association is required for both desmosome formation and maintenance. These studies identify the biophysical properties of desmoglein transmembrane domains as key determinants of lipid raft association and desmosome adhesive function.

## INTRODUCTION

The plasma membrane is a dynamic and heterogenous landscape of lipid, cholesterol, and protein that defines the cell boundary and integrates environmental cues with intracellular chemical and mechanical signaling pathways. In epithelial cells, intercellular junctions assemble into specialized plasma membrane domains that mediate cell adhesion and communication. The desmosome is a protein-dense and highly ordered intercellular junction that confers robust adhesion to prevent mechanically-induced tissue disruption (Skerrow and Matoltsy, 1974; Matoltsy, 1975; Hennings et al., 1980; Hennings and Holbrook, 1983; Bartle et al., 2017; Sikora et al., 2020). Within desmosomes, the desmosomal cadherins, desmogleins (DSG) and desmocollins mediate adhesion across the extracellular space while these proteins are intracellularly anchored to the intermediate filament cytoskeleton via plakoglobin, plakophilins, and desmoplakin (Gorbsky and Steinberg, 1981; Cowin et al., 1984; Kowalczyk et al., 1997; Bornslaeger et al., 2001). Numerous diseases are associated with faulty desmosome assembly (Zimmer and Kowalczyk, 2020), yet how desmosomal proteins form a densely packed, dynamic and functionally adhesive unit is poorly understood.

Lipid raft membrane domains are functional collections of lipids and membrane-associated proteins in an ordered lipid environment enriched in cholesterol, sphingolipids, and longer-chain saturated phospholipids (Simons and Ikonen, 1997; Levental and Lyman, 2023). Previous work demonstrated that desmosomes are a mesoscale lipid raft membrane domain (Lewis, Caldara et al., 2019) whose assembly and disassembly require lipid raft association of several desmosomal protein components (Nava et al., 2007; Resnik et al., 2011; Brennan et al., 2012; Jiang et al., 2014; Stahley et al., 2014; Resnik et al., 2019). Importantly, disease-causing point mutations in the DSG1 transmembrane domain (TMD) that reduce raft association also abrogate desmosome assembly and function (Lewis, Caldara et al., 2019; Zimmer et al., 202). However, the mechanisms through which desmosomal proteins associate with raft-like membrane domains are largely uncharacterized. Previous research has revealed the physical properties of TMDs that promote their association with raft phases (Levental et al., 2010; Diaz-Rohrer et al., 2014; Lorent et al., 2017). These properties include TMD length, TMD bulkiness, and TMD palmitoylation. Longer TMDs are more compatible with the thicker lipid bilayer of raft domains because they prevent hydrophobic mismatch between the TMD residues and phospholipids (Diaz-Rohrer et al., 2014). TMD bulkiness is determined by the size of amino acid side chains; TMDs composed of smaller residues are more amenable to the tightly packed and ordered lipid raft environment (Lorent et al 2017). Lastly, palmitoylation is a reversible post-translational modification that adds a raftophilic, saturated, 16-carbon acyl chain to cysteine residues on the intracellular end of TMDs (Melkonian et al., 1999; Levental et al., 2010). DSG1-4 possess highly conserved TMDs that are long, relatively non-bulky and contain palmitoylated TMD-adjacent cysteine residues (Roberts et al., 2016; Lewis, Caldara et al., 2019). The role of these TMD features in DSG raft association, desmosome assembly, and adhesive strength is unknown.

We hypothesized that DSG1 TMD physical properties, including length, bulkiness, and palmitoylation, drive DSG1 lipid raft association to promote desmosome assembly and function. We tested this hypothesis by expressing a panel of shortened, bulked, or palmitoylation-deficient DSG1_TMD_-GFP variants in a DSG- and desmosome-null background. We then used biochemical, imaging, and functional assays to assess whether DSG1_TMD_-GFP variants associate with rafts and assemble functional desmosomes. We found that reduced raft association strongly correlated with both desmosome assembly and adhesive strength. Though most DSG1_TMD_-GFP variants trafficked normally to the cell surface, altered dynamics at the plasma membrane resulted in reduced rates of desmosome assembly, increased surface turnover, or both. Overall, our findings indicate that TMDs govern desmoglein raft association to drive local clustering and nucleation of desmosomal proteins into stable adhesive complexes.

## RESULTS

### Palmitoylation does not affect DSG1 raft association

To determine the physical properties of the DSG1 TMD that govern raft association, we designed a panel of GFP-tagged DSG1_TMD_ variants to individually alter palmitoylation, TMD length, or hydrophobic surface area (bulkiness) (Table 1). The DSG1 TMD encompasses residues F546-I569. To prevent palmitoylation, we substituted juxtamembrane cysteine residues with alanine (DSG1_PALM_-GFP). To modify length, we truncated seven residues from either the N-terminal or C-terminal end of the DSG1_TMD_ (DSG1_Δ7N_-GFP; DSG1_Δ7C_-GFP) or we introduced a glycine to arginine substitution at G552 or G562 (DSG1_G552R_-GFP; DSG1_SAM_-GFP). To modify bulkiness, we substituted all TMD residues with leucines (DSG1_Leu_-GFP), inserted the non-raft associating TMD from the adherens junction protein, E-cadherin (DSG1_Ecad_-GFP), or inserted the raft-associating TMD from linker for activation of T-cells (LAT; DSG1_LAT_-GFP). Finally, we randomized (scrambled) the normal DSG1 TMD amino acid sequence (DSG1_scr_-GFP). We calculated the predicted raft partitioning free energy values, represented by ΔG_raft_, for each DSG1_TMD_ variant using a formula validated for a wide range of single-pass transmembrane proteins (Lorent et al., 2017) (Table 1). Since smaller ΔG_raft_ values indicate greater raft affinity, these calculations predicted DSG1_PALM_ to have the highest negative impact on raft affinity, suggesting that palmitoylation may drive DSG1 raft association.

**Table 1:**
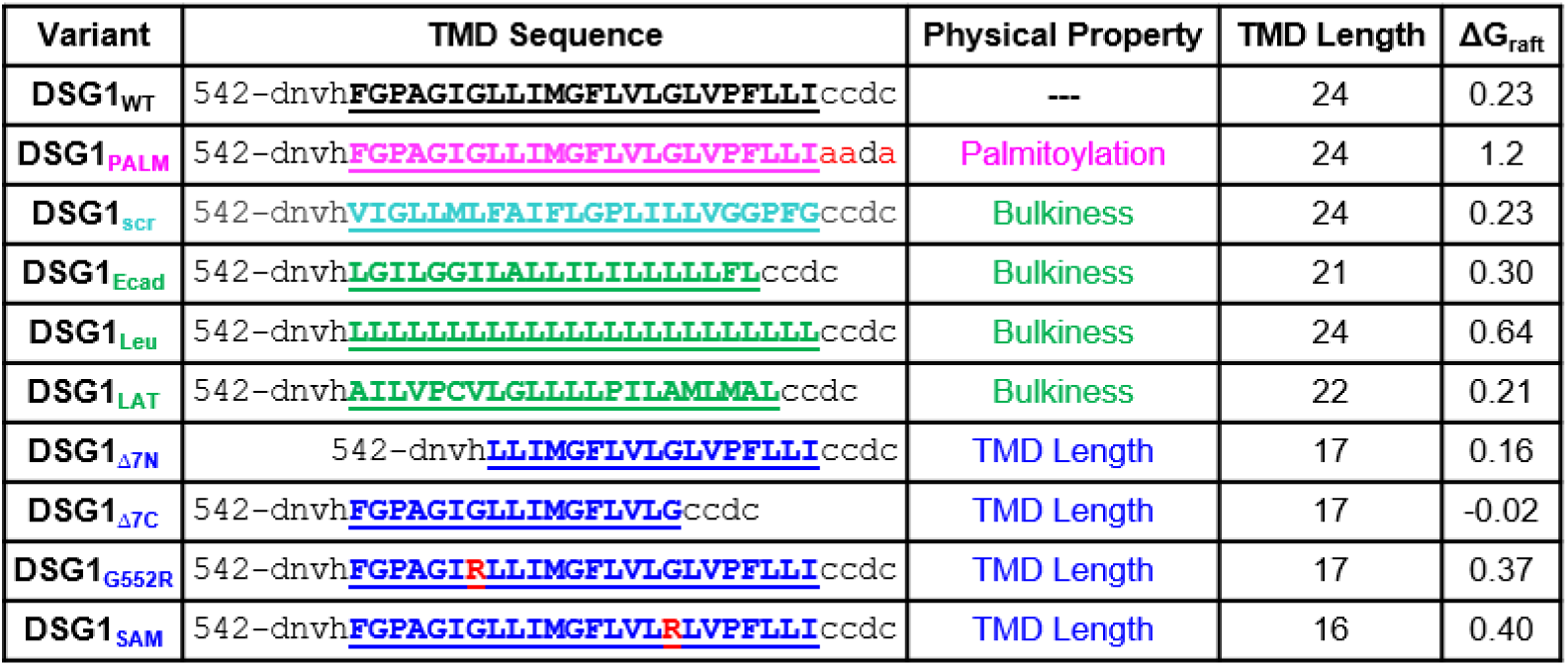
Sequence and properties of DSG1 TMD variants. DSG1 TMD variants alter physicochemical properties such as palmitoylation, exposed surface area, or TMD length. ΔG_raft_ calculations estimate raft association for each DSG1 TMD variant using an equation (Lorent et al., 2017).

To experimentally determine whether palmitoylation drives DSG1 raft association, we stably expressed DSG1_PALM_-GFP in DSG-null A431 cells. A431 cells are an immortal human carcinoma cell line commonly used to assess desmosome assembly and function (Godsel et al., 2005; Brennan et al., 2012; Baddam et al., 2018; Bharathan et al., 2023). DSG2 is the prevalent DSG isoform expressed in A431 cells (Schäfer et al., 1994, Zimmer et al., 2022). CRISPR-mediated knockout of DSG2 in A431 cells abrogates desmosome formation (Zimmer et al., 2022), allowing us to assess the ability of various DSG1 mutants to rescue desmosome assembly and function in a DSG-null background. Western blots of whole cell lysates from cells expressing DSG1_WT_-GFP or DSG1_PALM_-GFP revealed that these constructs were expressed at similar levels and that the expression levels of other desmosomal proteins including plakoglobin, desmocollin-2, and desmoplakin were unaffected (Supplemental Figure 1, A and E-H). To test the ability of DSG1_PALM_-GFP to associate with lipid rafts, we used sucrose gradient fractionations to separate detergent-resistant (raft) membranes (DRMs) from non-detergent-resistant membranes (non-DRMs) (Lingwood and Simons, 2007). DSG1_PALM_-GFP partitioned to DRMs to a similar extent as DSG1_WT_-GFP (Figure 1, A and B).

**Figure 1:**
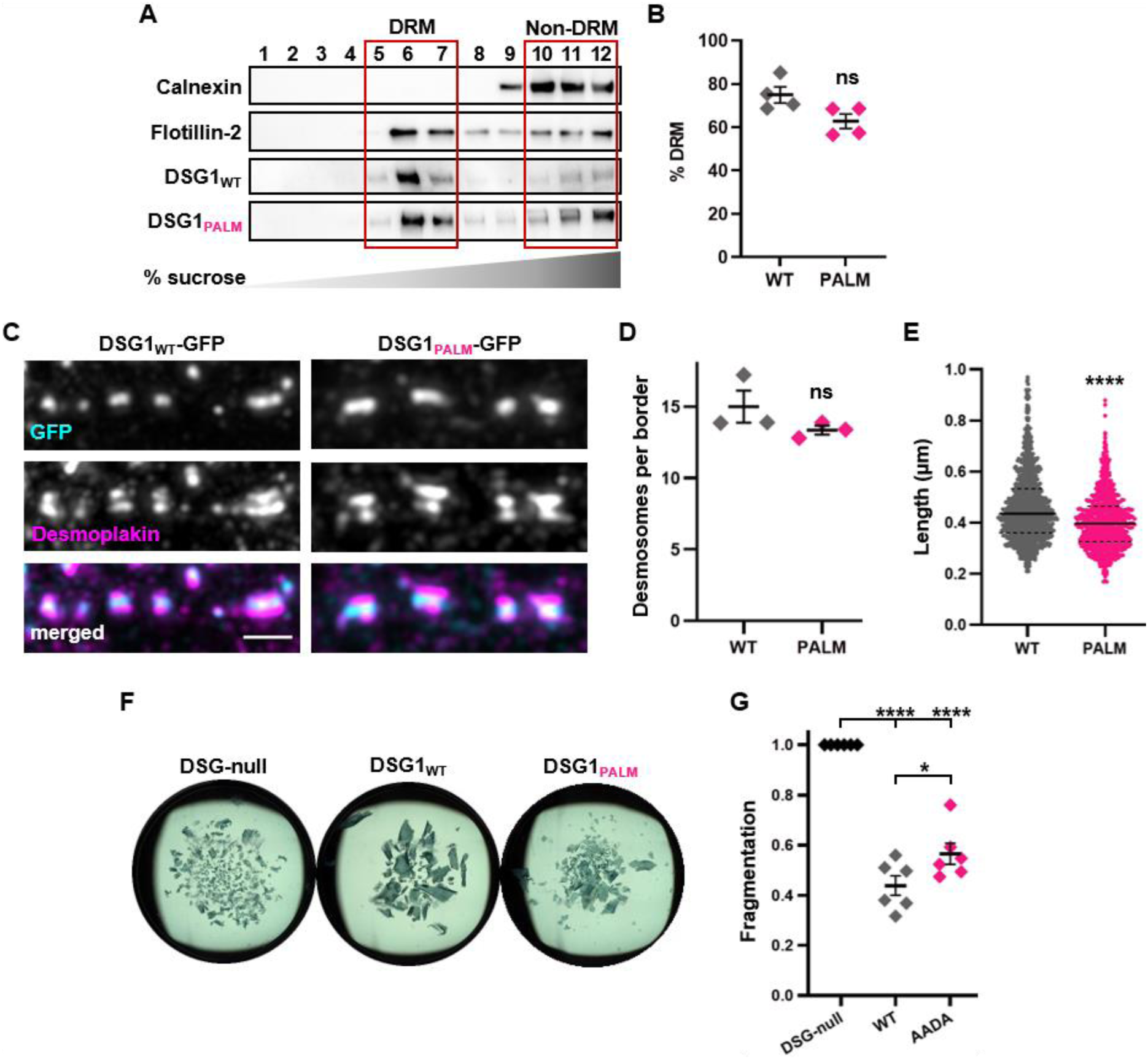
Palmitoylation-deficient DSG1 associates with rafts and assembles adhesive desmosomes. A) Sucrose gradient fractionations from DSG-null cells expressing DSG1_WT_-GFP or DSG1_PALM_-GFP show distribution of DSG1-GFP between DRM and non-DRM fractions. B) Quantification of westerns in (A), n = 4. C) Maximum intensity projections of SIM images from DSG-null cells expressing DSG1_WT_-GFP or DSG1_PALM_-GFP show desmosomes identified by desmoplakin railroad tracks. Bar, 1 µm. D-E) Quantification of desmosomes per border (D) and average desmosome length (µm) (E). Each point represents the average length of up to a 1000 measured desmosomes per replicate, n = 3. F) Dispase cell dissociation assay of DSG-null monolayers or monolayers from DSG-null cells expressing DSG1_WT_-GFP or DSG1_PALM_-GFP. Fewer fragments indicate strong desmosomal adhesion. G) Quantification of (F). Fragment counts normalized to monolayer fragmentation of DSG-null cells, n = 6.

Next, we asked whether DSG1_PALM_-GFP formed desmosomes. Desmoplakin ‘railroad tracks,’ an indicator of mature desmosomes, can be resolved with structured illumination microscopy (SIM) (Stahley et al., 2016). We found that cells expressing DSG1_PALM_-GFP formed apparently normal desmosomes (Figure 1, C). Defining desmosomes by the presence of DSG1-GFP sandwiched between desmoplakin ‘railroad tracks,’ we quantified both desmosome number and length. Desmosomes formed by DSG1_PALM_-GFP were of a similar number but shorter than those formed by DSG1_WT_-GFP (Figure 1, D and E). Finally, to determine whether desmosomes formed by DSG1_PALM_-GFP were functional, we used a dispase-based monolayer fragmentation assay in which confluent monolayers are lifted from their substrate and subjected to mechanical stress on an orbital shaker (Calautti et al., 1998; Huen et al., 2002). DSG-null cells fragment heavily in this assay due to a lack of desmosomes. Expression of DSG1_WT_-GFP rescues desmosome adhesion strength (Zimmer et al., 2022). Similarly, DSG1_PALM_-GFP also rescued desmosome function in these cells nearly as well as DSG1_WT_-GFP (Figure 1, F and G). These data indicate that palmitoylation is not required for DSG1 raft association or steady state desmosome adhesive function.

### TMD length and bulkiness determine DSG1 raft association

Since palmitoylation did not affect DSG1 raft association in our biochemical assay (Figure 1, A and B), we tested the effects of DSG1 TMD bulkiness or length on DSG1 raft association. As with DSG1_PALM_-GFP, we stably expressed these variants in DSG-null A431 cells. We used fluorescence-activated cell sorting to obtain cell populations expressing DSG1_TMD_-GFP variants at similar levels to DSG1_WT_-GFP. We observed no effects on the expression levels of other desmosomal proteins (Supplemental Figure 1).

Using sucrose gradient fractionations, we found that DSG1_WT_-GFP DRM partitioning in DSG-null cells is similar to DSG2 DRM partitioning in A431 cells (Figure 2, A and B). However, DSG1_short_ or DSG1_bulky_ variants displayed reduced DRM partitioning. Additionally, the desmosomal adaptor protein, plakoglobin, requires DSG for DRM partitioning as plakoglobin DRM partitioning is significantly decreased in DSG-null cells and fully rescued by expression of DSG1_WT_ (Supplemental Figure 2, A and B). In contrast, the expression of non-rafty DSG1_TMD_ variants is not sufficient to restore plakoglobin DRM partitioning. We observed a striking correlation (R^2^ = 0.882) between the DRM partitioning of DSG1_TMD_ variants and that of plakoglobin (Supplemental Figure 2, C). These findings suggest that TMD length and bulkiness drive raft association of DSG1 and that plakoglobin raft association depends on DSG1 raft association.

**Figure 2:**
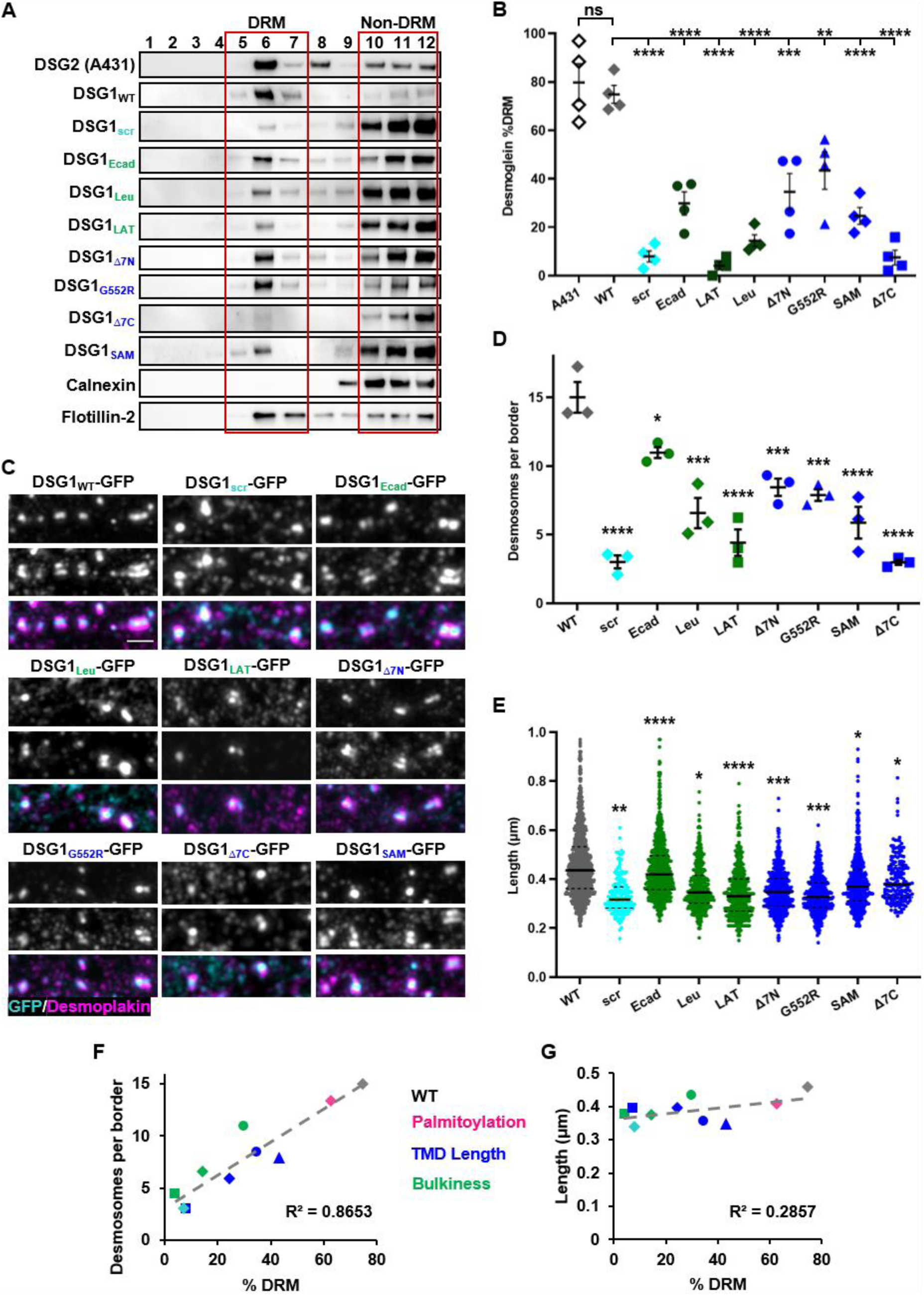
DSG1 raft association is determined by TMD length and bulkiness and supports desmosome formation. A) Sucrose gradient fractionations of DSG-null cells expressing DSG1_WT_-GFP or each of the DSG1_TMD_-GFP variants show distribution of DSG1-GFP between DRM and non-DRM fractions. B) Quantification of westerns in (A), n = 4. C) Maximum intensity projections of SIM images from DSG-null cells expressing DSG1_WT_-GFP or each of the DSG1_TMD_-GFP variants show desmosomes identified by desmoplakin railroad tracks. D) Quantification of desmosomes per border, n = 3. E) Quantification of desmosome length (µm). Each point represents the average length of up to a 1000 measured desmosomes per replicate, n = 3. F-G) Scatter plot of correlation between degree of raft association (% DRM) and desmosome number (F) or desmosome length (G), color-coded by TMD property. Point shape and color match those used in (B), (D), and (E) for identification.

### DSG1 raft association determines desmosome number and function

Using SIM to image cells stained for GFP and desmoplakin, we found that altering DSG1 TMD properties reduced desmosome number and size relative to DSG1_WT_-GFP (Figure 2, C-E). Importantly, DSG1_TMD_ variant raft association strongly correlated (R^2^ = 0.8653) with desmosome number (Figure 2, F). However, we found little correlation between DSG1 raft association and desmosome length (R^2^ = 0.2857) (Figure 2, G).

We also assessed the adhesive function of each DSG1_TMD_-GFP variant using the dispase-based monolayer fragmentation assay (Figure 3, A and B). We observed a striking correlation (R^2^ = 0.8551) between DSG1 raft association and adhesive strength (Figure 3, C). Further, we observed a strong correlation between desmosome number and adhesive strength (R^2^ = 0.6969), but little correlation between desmosome length and adhesion (R^2^ = 0.1636) (Figure 3, D and E). Given the strong relationship between the raft association of DSG1 and plakoglobin observed above, plakoglobin raft association similarly correlated strongly with desmosome number (R^2^ = 0.7058) and adhesive strength (R^2^ = 0.8441) but not with length (R^2^ = 0.2447) (Supplemental Figure 2, D-F). Altogether, these results suggest that DSG1_TMD_ length and bulkiness drive DSG1 and plakoglobin raft association to promote desmosome formation and function.

**Figure 3:**
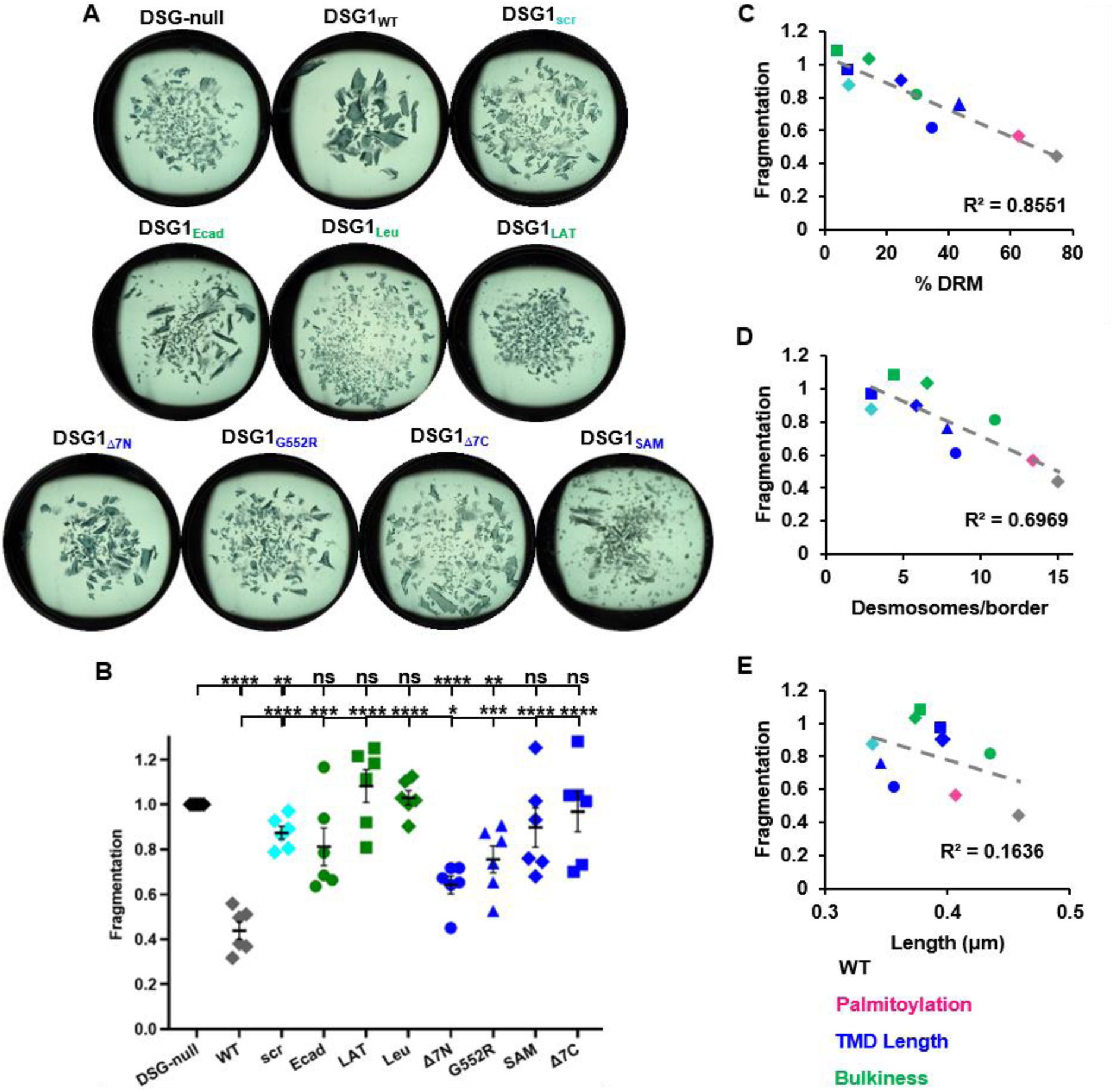
DSG1 raft association correlates with desmosome adhesive function. A) Dispase monolayer fragmentation assay of DSG-null monolayers or monolayers from DSG-null cells expressing DSG1_WT_-GFP or other DSG1_TMD_-GFP variants. Fewer fragments indicate strong desmosomal adhesion. B) Quantification of (A). Fragment counts normalized to monolayer fragmentation of DSG-null cells, n = 6. Statistically, all DSG1_TMD_ variants were compared to DSG-null monolayer fragmentation (top asterisks) to assess general ability to functionally rescue or to DSG1_WT_ (bottom asterisks) to assess function relative to WT conditions. C-E) Scatter plot of correlation between mean desmosome length (µm) in Figure 2E and fragmentation values from (B) (C), mean desmosome count in Figure 2D and fragmentation values from (B) (D), and % DRM in Figure 2B and fragmentation values from (B) (E).

### Reduced DSG1 raft association alters desmosome assembly dynamics

Having observed that our DSG1_TMD_-GFP variants represent a collective of raft affinities and adhesive functions at steady state, we sought to determine the relationships between raft association and desmosome number. We reasoned that one of three scenarios may be occurring in the presence of non-raft DSG1_TMD_-GFP variants: 1) desmosomes assemble more slowly, 2) desmosomes assemble at a normal rate but are unstable and disassemble quickly, or 3) desmosomes are both slow to assemble and quick to disassemble. We first compared desmosome assembly rates following a calcium switch. Cells were placed in low calcium medium for >18 hours to eliminate desmosomes, then back into high calcium medium for 1, 3, or 12 hours to activate new desmosome assembly. Cells were then fixed and imaged with a spinning disk confocal microscope (Figure 4, Supplemental Figure 3). We were able to distinguish desmoplakin ‘railroad tracks’ for counting and measuring at each timepoint (Figure 4, A; Supplemental Figure 3 and 4).

**Figure 4:**
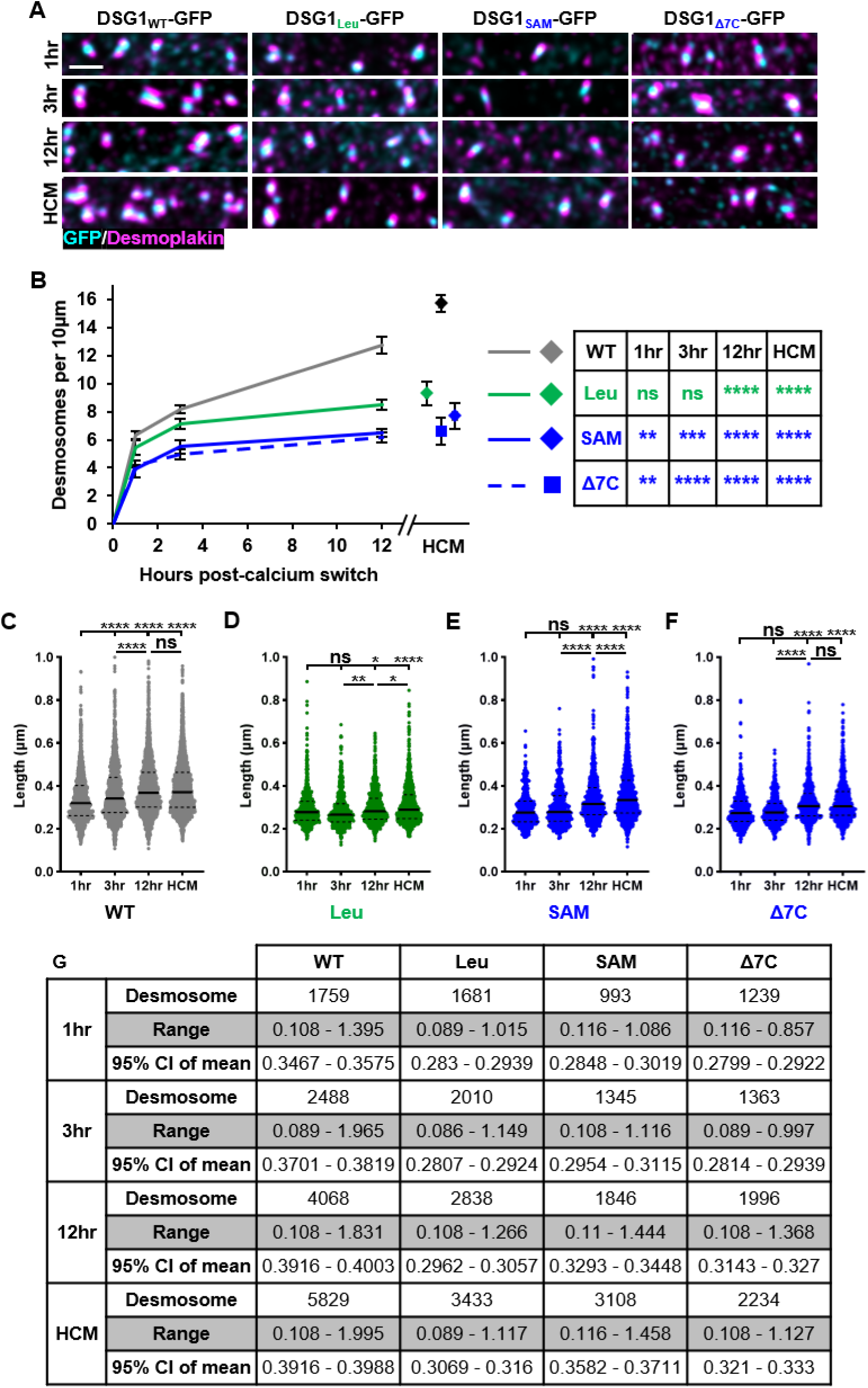
DSG1 raft association regulates desmosome dynamics. (A) Maximum intensity projections of spinning disk confocal images of DSG-null cells expressing DSG1_WT_-GFP, DSG1_Leu_-GFP, DSG1_SAM_-GFP, or DSG1_Δ7C_-GFP stained for GFP and DP at steady state or 1, 3, or 12 hours after a calcium switch. Bar, 1 µm. These variants are representative of three mechanistic categories into which the DSG1_TMD_ variants can be grouped. (B) Graph showing desmosome assembly progression following a calcium switch and compared to desmosome quantities at steady state. Error bars represent mean +/− SEM, n = 3. (C-F) Violin plots show distribution of measured desmosome lengths 1, 3, or 12 hours post-calcium switch or during steady state (HCM, high calcium medium) for DSG-null cells expressing DSG1_WT_-GFP (C), DSG1_Leu_-GFP (D), DSG1_SAM_-GFP (D), or DSG1_Δ7C_-GFP (F). Y-axis was limited to 1 µm to better visualize spreads. DSG1_WT_-GFP at 3hr, 12hr, and HCM and DSG1_SAM_-GFP at 12hr and HCM had limited measurements above 1 µm. G) Table describing total number of measured desmosomes (across three replicates), range of measured desmosome lengths (um), and 95% confidence interval (CI) of the mean after a 1hr, 3hr, or 12hr calcium switch or at steady state.

Cells expressing DSG1_WT_-GFP formed desmosomes rapidly upon calcium reintroduction and continued to steadily form desmosomes throughout the 12-hour time course (Figure 4 A and B; Supplemental Figure 3). The bulky DSG1_Leu_-GFP variant supported rapid desmosome growth at early time points, nearly matching DSG1_WT_-GFP up to 3 hours post calcium. However, desmosome formation in cells expressing DSG1_Leu_-GFP plateaued and failed to match desmosome numbers achieved by cells expressing DSG1_WT_-GFP. Similar behavior was observed for the DSG1_Ecad_-GFP, DSG1_Δ7N_-GFP, and DSG1_G552R_-GFP variants (Supplemental Figure 3). In contrast, cells expressing the DSG1_SAM_-GFP mutant exhibited fewer desmosomes at 1 hour, and desmosome number plateaued by 3 hours, with similar behaviors for DSG1_LAT_-GFP, DSG1_scr_-GFP, and DSG1_Δ7C_-GFP variants.

Desmosome length increases as junctions mature after a calcium switch (Beggs et al., 2022). We quantified the length of each counted desmosome and found that desmosomes in cells expressing DSG1_WT_-GFP increased in length over time with the longest desmosomes measured in high calcium steady state conditions (Figure 4, C and G; Supplemental Figure 4, A-E, Supplemental Table 1). Increases in desmosome length over time were also observed for cells expressing most of the DSG1_TMD_ variants (Figure 4, D-G; Supplemental Figure 4; Supplemental Table 1). However, these desmosomes were consistently smaller than those in cells expressing DSG1_WT_, and the observed changes in length were often inconsistent with fluctuations or initial increases followed by growth stagnation. Overall, the initial rate of desmosome formation, the length of desmosomes, and the number of desmosomes formed are influenced by DSG raft association through DSG TMD properties.

### DSG1 raft association regulates desmosome assembly and DSG1 surface turnover

Recent studies have linked raft association with the kinetics of secretory trafficking (Castello-Serrano et al., 2024), suggesting that the influence of raft affinity on desmosome assembly and steady-state characteristics may be related to DSG1 secretory kinetics. To test this hypothesis, we repeated the calcium switch experiments but included stains for GM130 and wheat germ agglutinin (WGA) to identify Golgi and plasma membrane pools of the DSG1_TMD_ variants, respectively (Figure 5; Supplemental Figure 5). DSG1_WT_-GFP and GM130 colocalization peaks 1-hour post-calcium switch while DSG1 colocalization with WGA increases rapidly after the first hour, plateauing at 3 hours. The other DSG1_TMD_-GFP variants exhibit similar behaviors except DSG1_scr_-GFP and DSG1_SAM_-GFP, which exhibit both delayed Golgi exit and plasma membrane localization. The other exception was DSG1_Δ7C_-GFP, which failed to appreciably localize to the plasma membrane. Despite delays for these variants, all DSG1_TMD_-GFP variants exhibit a low baseline level of colocalization with WGA in low calcium medium, indicating that a pool of DSG1 is present at the plasma membrane even in the absence of cell-cell contact. Though the delay in desmosome formation observed above (Figure 4; Supplemental Figure 3) for DSG1_SAM_-GFP, DSG1_scr_-GFP, and DSG1_Δ7C_-GFP variants can be explained in part by a failure to rapidly transit through the biosynthetic pathway, the presence of membrane-localized DSG1 without cell-cell contact suggests that the earliest desmosomes are likely formed from DSG1 already present at the plasma membrane.

**Figure 5:**
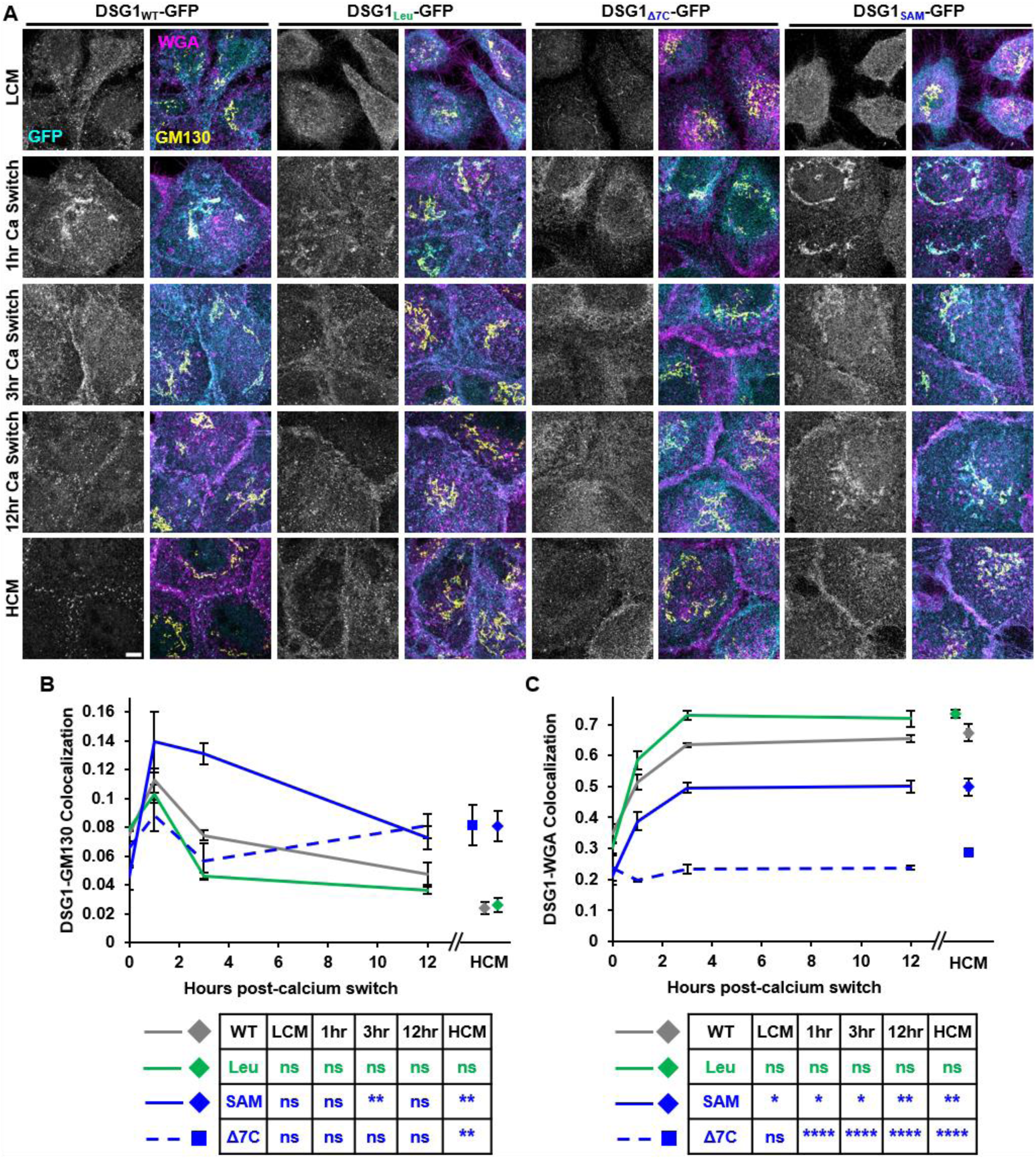
Most DSG1_TMD_ variants traffic normally through the Golgi body to the plasma membrane. A) Spinning disk confocal images show DSG-null cells expressing DSG1_WT_-GFP, DSG1_Leu_-GFP, DSG1_SAM_-GFP, or DSG1_Δ7C_-GFP and stained for GM130 and WGA to mark Golgi and plasma membranes, respectively. Bar, 5µm. B-C) Quantification of colocalization between DSG1_TMD_-GFP and GM130 or DSG1_TMD_-GFP and WGA. Error bars represent mean +/− SEM, n = 3.

Our analysis of desmosome assembly rates above (Figure 4) revealed that the number of desmosomes formed by several of the DSG1_TMD_-GFP variants plateaued earlier than DSG1_WT_-GFP. We hypothesized that these differences might reflect decreased cell surface stability. We assessed cell surface turnover rates by performing a pulse-chase experiment. Cell surface DSG1 was labeled with an antibody against the DSG1 extracellular domain. After labeling cell surface DSG1 at 4°C, cells were rinsed, moved to 37°C, fixed after chase without permeabilization, and then labeled with a fluorescent secondary antibody to exclusively assess cell surface DSG1 levels. We found that DSG1_PALM_-GFP, DSG1_scr_-GFP, DSG1_Leu_-GFP, DSG1_G552R_-GFP, DSG1_Δ7N_-GFP, and DSG1_Δ7C_-GFP exhibited increased surface turnover rates while DSG1_Ecad_-GFP, DSG1_LAT_-GFP, and DSG1_SAM_-GFP maintained similar surface turnover rates relative to DSG1_WT_-GFP (Figure 6, A and B, Supplemental Figure 6). We repeated this experiment using an antibody against the desmocollin-2 extracellular domain and found similar patterns such that desmocollin-2 showed faster surface turnover in the presence of DSG1_TMD_-GFP variants that exhibit increased surface turnover (Figure 6, C-E; Supplemental Figure 7).

**Figure 6:**
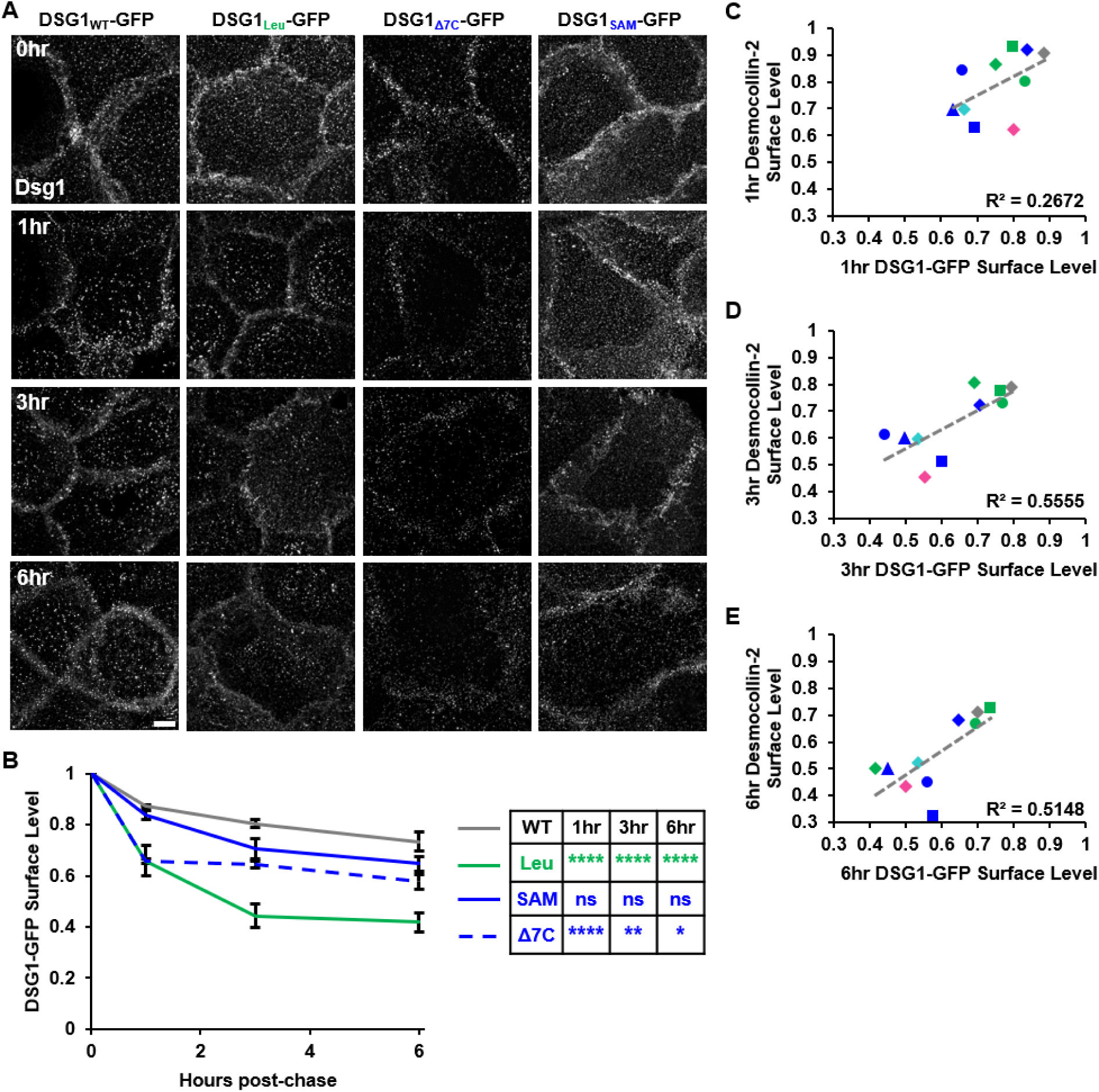
Many DSG1_TMD_-GFP variants exhibit increased surface turnover and increased desmocollin-2 surface turnover. A) Maximum intensity projections of images show surface level DSG1 in DSG-null cells expressing DSG1_WT_-GFP, DSG1_Leu_-GFP, DSG1_SAM_-GFP, or DSG1_Δ7C_-GFP after 0, 1, 3, or 6 hours of turnover. Bar, 5 µm. B) Quantification of images in (A). Error bars represent mean +/− SEM, n = 3. C-E) Scatterplot showing correlation between surface levels of DSG1 or desmocollin-2 after 1-hour (C), 3-hour (D), or 6-hour (E) pulse-chase normalized to 0 hour. Desmocollin-2 surface turnover results are in Supplemental Figure 7.

## DISCUSSION

Desmosomal proteins utilize several clustering mechanisms to form an interlocking assembly of interactions where extracellular adhesion and intracellular adaptor proteins anchor desmosomal cadherins into a dense, ordered, mechanically-resistant and adhesive protein complex. These clustering mechanisms include adhesion mediated by the extracellular domains of DSG and desmocollin in both *cis* and *trans* (Chitaev and Troyanovsky, 1997; Wu et al., 2010; Nie et al., 2011; Al-Amoudi et al., 2011; Harrison et al., 2016) and clustering promoted by intracellular adaptor proteins (Kowalczyk et al., 1997; Fuchs et al., 2019, Zimmer et al., 2020; Wanuske et al., 2021). Here, we report that the intrinsic propensity for the desmosomal cadherin TMDs to segregate into lipid raft domains provides an additional clustering mechanism. Specifically, lipid raft affinity of the DSG1 TMD is driven by unique properties of the TMD, and reducing raft affinity by altering these TMD properties is sufficient to impede desmosome assembly and function. This intrinsic propensity helps to explain why disease-causing DSG1 mutations that abrogate raft association cause desmosomal defects and epidermal fragility (Lewis, Caldara, et al., 2019; Zimmer et al., 2022). Furthermore, these findings advance our understanding of the biophysical properties of the DSG1 TMD in raft association and further demonstrate the significance of this domain in desmoglein function.

Our panel of DSG1_TMD_ variants reveals that modulating the physical TMD properties of length and bulkiness consistently reduces DSG1 raft association while palmitoylation is not required for DSG1 raft association (Figures 1-2). The lack of a role for palmitoylation in DSG1 raft association is consistent with previous reports that palmitoylation-deficient DSG2 and DSG3 retain raft association (Roberts et al., 2016). TMD length and bulkiness were previously posited as drivers of DSG1 raft association in relation to two different disease-causing point mutations in the DSG1 TMD (Zimmer et al., 2022). Given the thickened lipid bilayer of desmosomes relative to non-desmosomal bilayers (Lewis, Caldara et al., 2019), a shorter, bulkier DSG TMD is energetically unfavorable for desmosome incorporation. In line with this reasoning, we consistently observe impaired desmosome formation and adhesion from each DSG1_SHORT_-GFP or DSG1_BULKY_-GFP variant (Figures 1-4; Supplemental Figures 3 and 4). Our results further indicate that the formation of raft domains by DSG1 is an intrinsic property of the protein and is coupled to DSG1 function. Importantly, we used a DSG-null system where DSG1_WT_-GFP fully rescues the desmosome-deficient phenotype whereas non-raft DSG1_TMD_-GFP variants are consistently insufficient. Therefore, DSG1 raft association, driven by TMD physical properties, is necessary to support normal desmosome formation and adhesion.

To determine the relationship between DSG1 raft association and desmosome formation, we explored the dynamics of DSG1 trafficking, desmosome assembly rates, and DSG1 surface turnover. Raft affinity promotes Golgi exit kinetics with non-raft proteins exiting at a slower rate than raft proteins (Castello-Serrano et al., 2024), but only a few of our non-raft DSG1_TMD_-GFP variants exhibit reduced trafficking rates. All DSG1_TMD_-GFP variants are present at the cell surface even in low calcium. Likewise, cell surface levels of the variants increase upon cell-cell contact induction (Figure 5; Supplemental Figure 5). Furthermore, only a few of the non-raft DSG1_TMD_-GFP variants exhibit reduced desmosome assembly rates during the first few hours after calcium-induced desmosome assembly, while the rates of desmosome assembly plateau at later timepoints for nearly all non-raft DSG1_TMD_-GFP variants (Figure 4; Supplemental Figure 3). Finally, the observed desmosome assembly rate plateaus are likely due to increased DSG1 and DSC2 surface turnover, which we observe for many but not all of the DSG1_TMD_-GFP variants (Figure 6; Supplemental Figures 6 and 7). Overall, each non-raft DSG1_TMD_-GFP variant forms fewer, poorly adhesive desmosomes through differing mechanisms involving combinations of reduced DSG1 trafficking rates, reduced desmosome assembly rates and/or increased DSG1 surface turnover rates (Figure 7). We conclude that DSG1 raft association promotes stability at the plasma membrane which is essential for desmosome maintenance and function; however, DSG1 surface stability is not sufficient for desmosome maintenance and function without adequate raft association.

**Figure 7:**
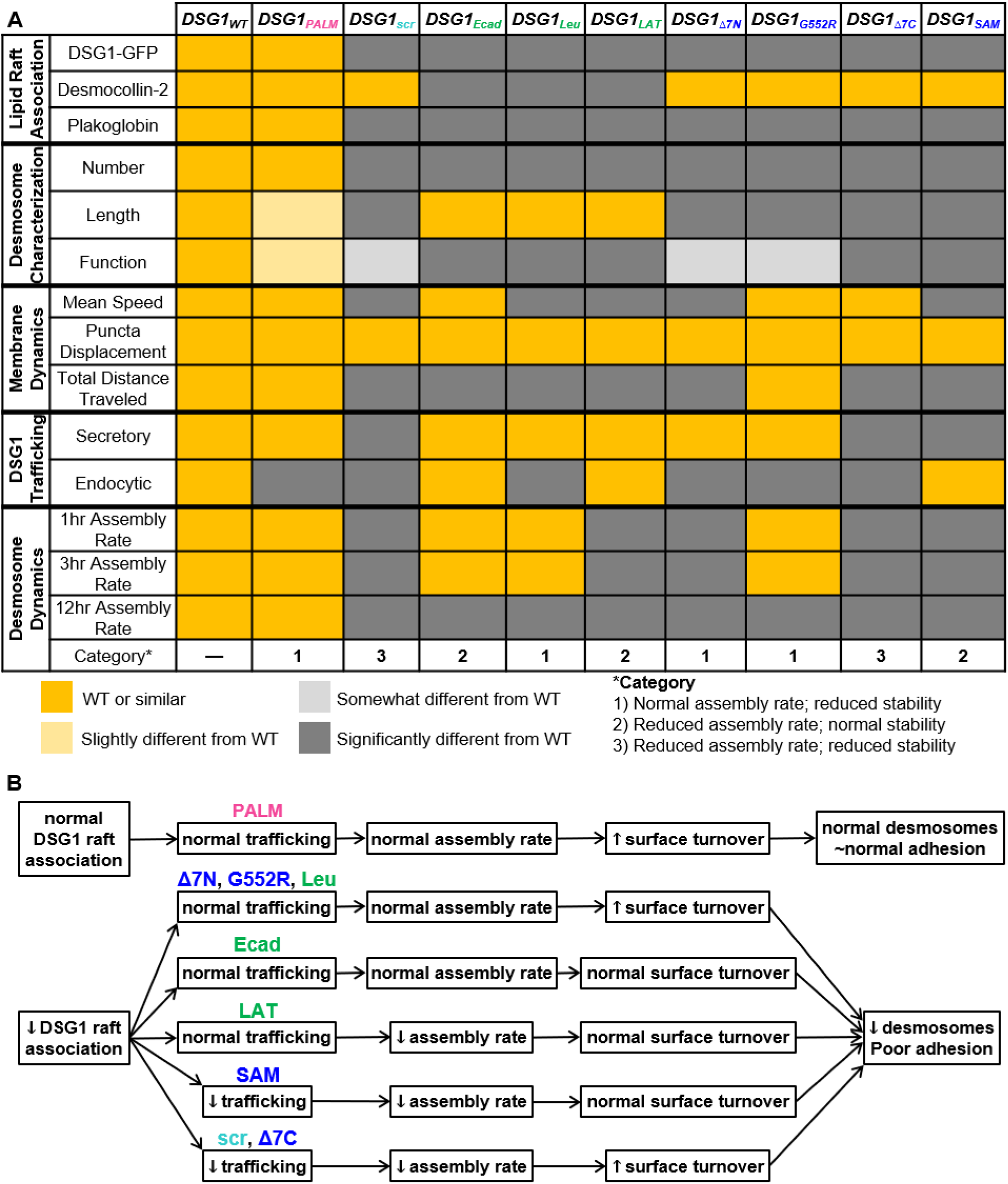
DSG1_TMD_ variant results summary. A) Color-coded tabular results summary illustrates how DSG1_TMD_ variants behave relative to DSG1_WT_ throughout the experiments performed in this work. B) The non-raft Dsg1_TMD_ variants consistently form fewer, poorly adhesive desmosomes through inconsistent pathways involving differing combinations of reduced trafficking rate, reduced desmosome assembly rate, and/or increased surface turnover.

Overall, our findings indicate that a pool of DSG1 present at the cell surface in low calcium conditions initiates desmosome formation upon a low-to-high calcium switch, followed by contributions from a biosynthetic pool that drives further desmosome formation and expansion (Figures 4-5; Supplemental Figures 3-5). In these conditions, DSG1 recruits desmosomal proteins to stably cluster by nucleating a lipid raft environment (Figure 8). DSG1 TMD length, bulkiness and palmitoylation then work in concert to promote desmosome assembly and regulate maintenance (Figures 1-2, 4, 6 and Supplemental Figures 2-4, 6, and 7).

**Figure 8:**
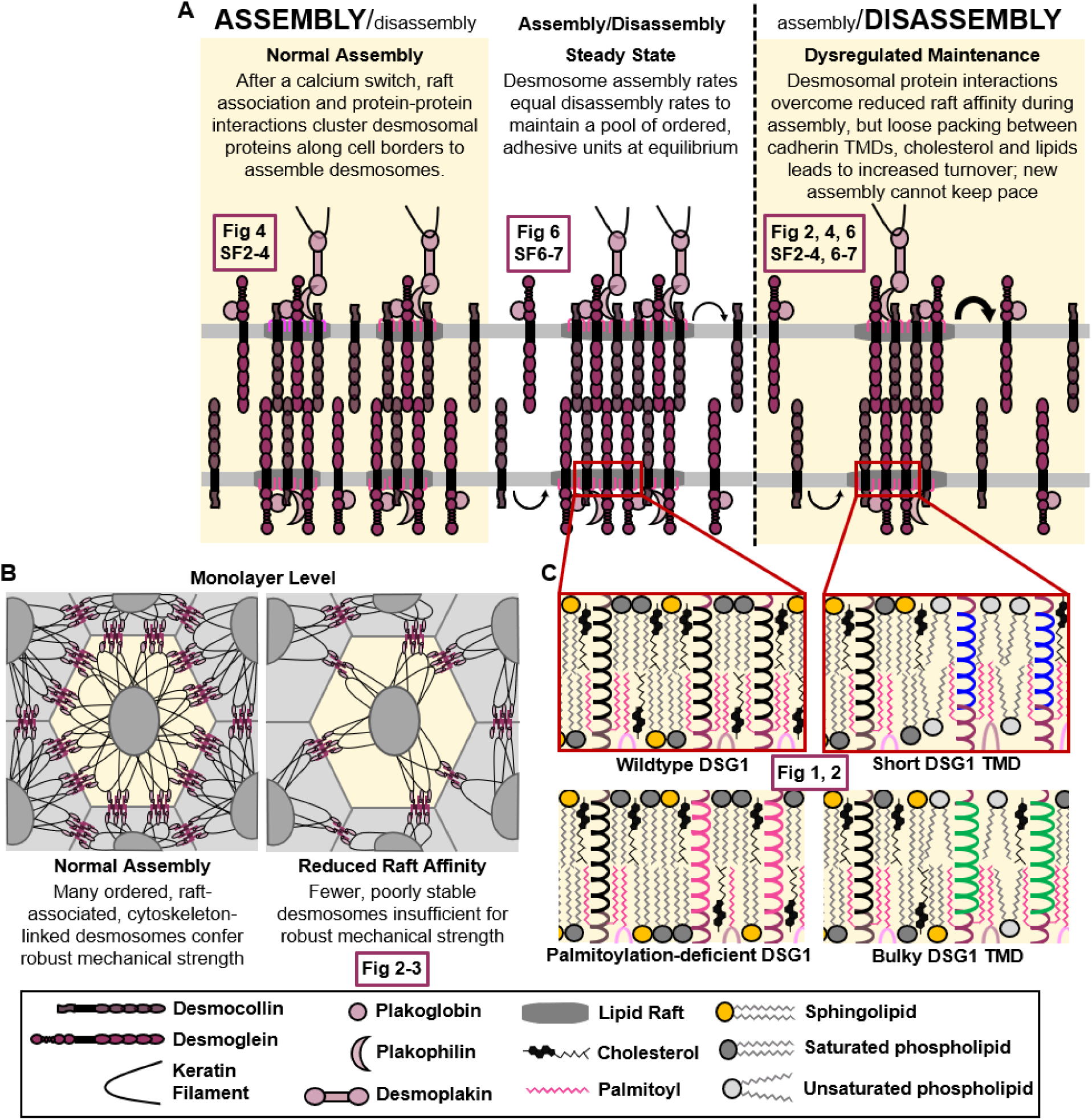
Schematic representation of desmosome maintenance dynamics. A) Addition of calcium to non-adhesive cells drives desmosome assembly by clustering desmosomal proteins via protein-protein interactions and raft association. This assembly-driven environment persists until cells have a pool of ordered, adhesive desmosomes at which point assembly and disassembly equilibrate to maintain a stable and adhesive population at steady state. Reduced DSG raft association alters this dynamic such that protein-protein interactions force the assembly of non-stable desmosomes. Cells attain desmosomal steady state faster due to increased DSG and DSC turnover that favors disassembly. B) Monolayer adhesion in the presence of raft-associated DSG1 versus non-raft associated DSG1: fewer, poorly adhesive desmosomes are insufficient for robust mechanical strength. C) Zoomed-in representation of the arrangement of palmitoylated desmosomal proteins, cholesterol, sphingolipid, and saturated phospholipids in the presence of wildtype DSG1 versus palmitoylation-deficient, short, or bulky DSG1. Shortened or bulked DSG1 TMDs disrupt lipid order. Boxes indicate the figure containing supporting relevant data.

The work presented here advances our understanding of the molecular interplay of DSG1 and the plasma membrane bilayer to identify features of the DSG1 TMD essential for desmosome assembly, disassembly, and adhesive function. Additional studies are needed to determine how the DSG TMD engages in TMD-protein, TMD-lipid, or TMD-cholesterol interactions that may promote desmosome assembly. Additionally, this work leaves open several questions about the relationship between raft association and other desmosomal functions, such as the regulation of signaling pathways associated with epidermal differentiation (Nekrasova et al., 2018; Cohen-Barak et al., 2020; Broussard et al., 2021).

## MATERIALS AND METHODS

### Generation of mutants and lentivirus

Cloning of plasmids expressing murine DSG1a_WT_-GFP and DSG1a_SAM_-GFP was previously described. For DSG1_LAT_-GFP, DSG1_G552R_-GFP, DSG1_Ecad_-GFP, and DSG1_Δ7N_-GFP, short sequences bearing the DSG1_TMD_ sequence variant flanked by appropriate restriction sites were obtained from Invitrogen’s Gene Art Strings and inserted into the plasmid expressing DSG1_WT_-GFP using existing restriction sites. Sequencing was performed through Genewiz to verify clones. The plasmids expressing DSG1_Leu_-GFP, DSG1_scr_-GFP, DSG1_PALM_-GFP, and DSG1_Δ7C_-GFP were ordered through Genscript. Lentivirus for DSG1_WT_-GFP and DSG1_SAM_-GFP was purchased from Cyagen VectorBuilder. Lentivirus for all remaining plasmids was made by co-transfection into HEK-293T cells together with pMD2.G (encoding VSV-G) and psPAX2 (encoding Gag and Pol). Lentivirus was collected from culture supernatants 24, 48, and 72 hours after transfection and concentrated by high-speed centrifugation.

### Cell line generation, culture, and reagents

Established A431 DSG-null cells (Zimmer et al., 2022) were cultured in DMEM (Corning; 4.5 g/L glucose, L-glutamine, and sodium pyruvate) with 10% fetal bovine serum (Hyclone) and 1X antibiotic-antimycotic solution (Corning). DSG-null cells were stably infected with lentiviruses expressing generated DSG1_TMD_-GFP variant constructs followed by selection with 5 μg/ml blasticidin (Thermofisher Scientific). No clonal isolation was performed. Cell lines expressing DSG1_TMD_-GFP variants were subjected to FACS with equivalent gating in order to remove highly-expressing and non-expressing cells to obtain populations with roughly equal DSG1_TMD_-GFP expression levels.

### Isolation of DRM

DRMs were isolated as described previously (Lingwood and Simons, 2007). Briefly, cells were cultured in 10-cm^2^ dishes and washed with PBS+. Cells were collected by scraping in TNE buffer containing protease inhibitors (Roche) and pelleted by centrifugation at 0.4xg at 4°C for 5 min (5415R; Eppendorf). Cells were resuspended in TNE buffer and homogenized using a 25-gauge needle. TNE buffer containing 1% Triton X-100 was added and cells were incubated on ice for 30 min. Detergent extract (420µm) was mixed with 840μl of 56% sucrose in TNE and placed at the bottom of a centrifuge tube. Volumes (1.9μl) of 35% and 5% sucrose were layered on top of the sample. Following an 18-hr centrifugation at 4°C (44,000 rpm, SW55 rotor, Beckman Optima LE-80 K Ultracentrifuge), 420-μl fractions (1-11, remaining volume combined to make up fraction 12) were removed from top to bottom of the gradient and stored at −20°C until processed for western blot analysis. Antibodies against flotillin-2 and calnexin were used as DRM and non-DRM markers, respectively.

### Immunofluorescence

Cells were cultured to 70% confluence on #1.5 glass coverslips. For experiments involving calcium switches, cells were maintained in low calcium medium (calcium-free, glutamine-free DMEM with 10% calcium-free FBS, 200 mM L-glutamine, 30 uM CaCl_2_) until 70% confluent and then subjected to calcium switch for 1 hour, 3 hours, or 12 hours before fixation.

For SIM and desmosome assembly rate experiments, cells were fixed in 3.7% paraformaldehyde (PFA) in PBS+ at 4°C for 12 min, rinsed in PBS+ containing 0.05% Triton X-100, and then blocked and permeabilized in PBS+ containing 0.1% Triton X-100 and 3% BSA for 30 min. Primary and secondary antibodies (listed below) were diluted into blocking solution (PBS+ containing 3% BSA and 0.05% Triton X-100). Coverslips were mounted to glass microscope slides using Prolong Gold (SIM) or Prolong Diamond (desmosome assembly rate) mounting medium (ThermoFisher Scientific).

For intracellular trafficking experiments, plasma membranes were labeled with WGA-568 (1 μg/ml; Biotium) for 15 min prior to fixation in 2% PFA for 15 min at 37°C before rinsing in PBS+ containing 0.05% saponin and then blocked and permeabilized in PBS+ containing 0.1% BSA and 0.05% saponin. Primary antibodies against GM130 and GFP and secondary antibodies (listed below) were diluted into blocking solution (PBS+ containing 1% BSA and 0.05% saponin). Coverslips were mounted to glass microscope slides using Prolong Diamond mounting medium (ThermoFisher Scientific).

For surface level turnover experiments, cells were pulsed for 30 min with primary antibodies against DSG1 or desmocollin-2 extracellular domains (listed below) and then chased with normal media for 0-hour, 1-hour, 3-hours, or 6-hours before fixation in 3.7% PFA. All buffers were the same as those used above except that Triton X-100 was excluded to prevent permeabilization and ensure that only surface levels would be labeled with the secondary antibody. Coverslips were mounted to glass microscope slides using Prolong Diamond mounting medium (ThermoFisher Scientific).

### Antibodies

The following primary antibodies were used: chicken-anti-GFP (1:1000, Aves 1020, immunofluorescence), rabbit-anti-GFP (1:1000, Invitrogen A11122, western blot), mouse-anti-GFP (1:1000, Abcam A1218, immunofluorescence), rabbit-anti-desmoplakin (1:1000, Bethyl A303-356A, immunofluorescence, western blot), mouse-anti-plakoglobin (1:1000, BD Transduction Laboratories 610254, western blot), mouse-anti-desmocollin-2/3 7G6 (1:250, Invitrogen 32-6200, immunofluorescence, western blot), human-anti-DSG1 PF1-8-15 (1:1500, gift from Aimee Payne, immunofluorescence), rabbit-anti-calnexin (1:500, Enzo ADI-SPA-86C-F, western blot), mouse-anti-flotillin-2 (1:1000, BD Transduction Laboratories 610383, western blot), mouse-anti-E-cadherin (1:500, Abcam 1416, immunofluorescence), mouse-anti-GM130 (1:1000, BD Transduction Laboratories 610822, immunofluorescence), mouse-anti-β-actin (1:3000, Sigma A1978, western blot). Secondary antibodies conjugated to Alexa Fluors (1:1000) were purchased from Invitrogen. Horseradish peroxidase-conjugated secondary antibodies (1:3000) were purchased from BioRad.

### Dispase-based monolayer fragmentation assay

Cells were cultured until confluent in 24-well tissue culture plates and treated with 1 U/ml dispase (Corning) for 15 min at 37°C. Cell sheets released from the plate were rinsed with PBS+, transferred to 1.5 ml Eppendorf tubes in 500 μl PBS+, and then subjected to mechanical stress on an orbital shaker on its highest setting for 45 sec. Fragments were transferred to a fresh 24-well plate, fixed with 1% PFA, and stained with methylene blue (Sigma). Plates were imaged on an Elispot scanner (Cellular Technologies, Ltd), and fragments were manually counted in Fiji (Schindelin et al., 2012).

### Image acquisition

SIM was performed using a Nikon N-SIM system on an Eclipse Ti-E microscope equipped with a 100x/1.49 NA oil immersion objective, 488- and 561-nm solid-state lasers in three-dimensional SIM mode. Images were captured using an EM charge-couple device camera (DU-897; Andor Technology) and reconstructed using NIS-Elements software with the N-SIM module (version 5.02; Nikon). Spinning disk confocal microscopy was performed using a Nikon Ti2-E equipped with a Yokogawa CSU-XI spinning disk unit, LUNF XL laser unit, Nikon Perfect Focus System, Z piezo stage, motorized XY stage, sCMOS camera (ORCA-Fusion BT, Hamamatsu Corp.). A Nikon 100x/1.49 NA Apo TIRF oil immersion objective was used with the correction collar positioned for room temperature imaging. For analysis of desmosome assembly rates, an additional 1.5x intermediate magnification was used to achieve sufficient resolution for resolving desmoplakin railroad tracks.

### Image Analysis and Statistics

Figure images are maximum intensity projections. All image processing and analysis was performed using Fiji (Schindelin et al., 2012). Images were deconvolved using Microvolution (Bruce and Butte, 2013). For studies quantifying desmosome number and length, desmosomes were measured using Straight lines in Fiji and collected in ROI manager for counting. For colocalization studies, channels were aligned using a mask created from images of fluorescent beads with NanoJ (Laine et al., 2019). Overlap between DSG1_TMD_-GFP and GM130 or DSG1_TMD_-GFP and WGA-568 was analyzed with BIOP JACoP (Bolte and Cordelières, 2006) using the Triangle (DSG1_TMD_-GFP; Zack et al., 1977) and Otsu (WGA-568; Otsu, 1979) thresholds or a manual threshold of 700 for GM130. For surface turnover studies, the integrated density of individual DSG1-GFP or DSC2 puncta was calculated using 3D Object Counter v2.0 (Bolte and Cordelières, 2006). Integrated densities were summed per image to represent total surface levels of DSG1 or DSC2 at each timepoint and averaged across 10 images per timepoint. Later timepoints were normalized to 0hr to calculate DSG1 or DSC2 surface level loss over time.

Statistics were calculated using Graphpad Prism 10.2.1. Error bars represent standard error of the mean. For most experiments, significance was determined using one-way ANOVA followed by Dunnett’s post-hoc and p-values have been indicated (ns, not significant, *p < 0.05, **p < 0.01, ***p < 0.001, ****p < 0.0001). For statistical analysis of desmosome length over time (Figure 4, C-G and Supplemental Figure 4), two-way ANOVA followed by Šídák’s multiple comparisons test was performed using the following length comparisons: 1-hour versus 3-hour, 1-hour versus 12-hour, 1-hour versus HCM, 3-hour versus 12hour, and 12h-hour versus HCM. Statistical analysis of immunofluorescence results was conducted on three independent experiments with 10 images per condition per replicate. For sucrose gradient fractionation, statistical analysis was conducted on results from four independent experiments. For expression level analysis on whole cell lysates, statistical analysis was conducted on results from three independent experiments. Statistical analysis of dispase assays was conducted on results from six independent experiments.

## ACKNOWLEDGEMENTS

The authors thank Aimee Payne for the use of a DSG1 antibody. The authors thank Dr. Navaneetha Bharathan for helpful manuscript review. This work was supported by National Institute of Health grants R01AR048266 and R01AR081883. SEZ was supported by the NIH training grant T32GM008367. This research project was also supported by the Emory University Integrated Cellular Imaging Microscopy Core (RRID:SCR_0234534) and the Emory Flow Cytometry Core.

