## Supplemental Figures for "The transmembrane domain of the desmosomal cadherin desmoglein-1 governs lipid raft association to promote desmosome adhesive strength"

**Supplemental Table 1**

| | | WT | PALM | scr | Ecad | LAT | Leu | Δ7N | G552R | SAM | Δ7C |
|---|---|---|---|---|---|---|---|---|---|---|---|
| 1hr | R1 | 582 | 589 | 560 | 530 | 354 | 687 | 515 | 557 | 368 | 542 |
| | R2 | 685 | 504 | 381 | 584 | 349 | 545 | 397 | 537 | 358 | 360 |
| | R3 | 492 | 484 | 274 | 636 | 370 | 449 | 483 | 403 | 267 | 337 |
| | **Total** | **1759** | **1577** | **1215** | **1750** | **1073** | **1681** | **1395** | **1497** | **993** | **1239** |
| | Range | 0.108 | 0.108 | 0.089 | 0.108 | 0.116 | 0.089 | 0.068 | 0.108 | 0.116 | 0.116 |
| | | **1.395** | **1.258** | **0.998** | **1.218** | **1.195** | **1.015** | **0.99** | **1.381** | **1.086** | **0.857** |
| | 95% CI of mean | 0.3467 | 0.2994 | 0.2735 | 0.2938 | 0.2906 | 0.283 | 0.2781 | 0.319 | 0.2848 | 0.2799 |
| | | **0.3575** | **0.313** | **0.2865** | **0.307** | **0.3073** | **0.2939** | **0.2906** | **0.3348** | **0.3019** | **0.2922** |
| 3hr | R1 | 885 | 833 | 778 | 713 | 505 | 746 | 474 | 767 | 469 | 479 |
| | R2 | 843 | 680 | 619 | 762 | 400 | 620 | 738 | 635 | 391 | 428 |
| | R3 | 760 | 636 | 435 | 674 | 542 | 644 | 631 | 578 | 485 | 456 |
| | **Total** | **2488** | **2149** | **1832** | **2149** | **1447** | **2010** | **1843** | **1980** | **1345** | **1363** |
| | Range | 0.089 | 0.108 | 0.089 | 0.108 | 0.108 | 0.086 | 0.108 | 0.096 | 0.108 | 0.089 |
| | | **1.965** | **1.469** | **0.973** | **1.294** | **1.063** | **1.149** | **1.076** | **1.096** | **1.116** | **0.997** |
| | 95% CI of mean | 0.3701 | 0.3527 | 0.2823 | 0.3258 | 0.3146 | 0.2807 | 0.2652 | 0.316 | 0.2954 | 0.2814 |
| | | **0.3819** | **0.3685** | **0.2927** | **0.3404** | **0.3303** | **0.2924** | **0.2748** | **0.3305** | **0.3115** | **0.2939** |
| 12hr | R1 | 1970 | 1432 | 1138 | 1041 | 698 | 969 | 903 | 994 | 574 | 747 |
| | R2 | 1151 | 1553 | 1017 | 859 | 670 | 974 | 929 | 681 | 571 | 677 |
| | R3 | 947 | 1361 | 669 | 1105 | 691 | 895 | 912 | 1013 | 701 | 572 |
| | **Total** | **4068** | **4337** | **2824** | **3005** | **2059** | **2838** | **2744** | **2688** | **1846** | **1996** |
| | Range | 0.108 | 0.125 | 0.108 | 0.086 | 0.108 | 0.108 | 0.089 | 0.091 | 0.11 | 0.108 |
| | | **1.831** | **1.913** | **1.352** | **1.511** | **1.218** | **1.266** | **1.163** | **1.574** | **1.444** | **1.368** |
| | 95% CI of mean | 0.3916 | 0.3516 | 0.3428 | 0.3399 | 0.3365 | 0.2962 | 0.2888 | 0.3429 | 0.3293 | 0.3143 |
| | | **0.4003** | **0.3605** | **0.3543** | **0.3522** | **0.3508** | **0.3057** | **0.2981** | **0.356** | **0.3448** | **0.327** |
| HCM | R1 | 1829 | 1656 | 900 | 2062 | 726 | 738 | 1024 | 1197 | 1272 | 559 |
| | R2 | 2115 | 1851 | 1154 | 1981 | 727 | 1399 | 1134 | 1272 | 1029 | 831 |
| | R3 | 1885 | 2168 | 929 | 1614 | 997 | 1296 | 1091 | 1462 | 807 | 844 |
| | **Total** | **5829** | **5675** | **2983** | **5657** | **2450** | **3433** | **3248** | **3931** | **3108** | **2234** |
| | Range | 0.108 | 0.125 | 0.086 | 0.122 | 0.086 | 0.089 | 0.086 | 0.108 | 0.116 | 0.108 |
| | | **1.995** | **1.684** | **1.545** | **1.531** | **1.355** | **1.117** | **1.077** | **1.634** | **1.458** | **1.127** |
| | 95% CI of mean | 0.3916 | 0.349 | 0.3073 | 0.3667 | 0.3442 | 0.3069 | 0.295 | 0.3699 | 0.3582 | 0.321 |
| | | **0.3988** | **0.3569** | **0.3171** | **0.3764** | **0.3583** | **0.316** | **0.3037** | **0.3817** | **0.3711** | **0.333** |

**Supplemental Table 1:** Number of desmosomes counted and measured (per repeat and totaled), range of measured lengths, and 95% confidence interval (CI) of the mean represented in violin plots in Figure 4 and Supplemental Figure 4.

### Supplemental Figure 1

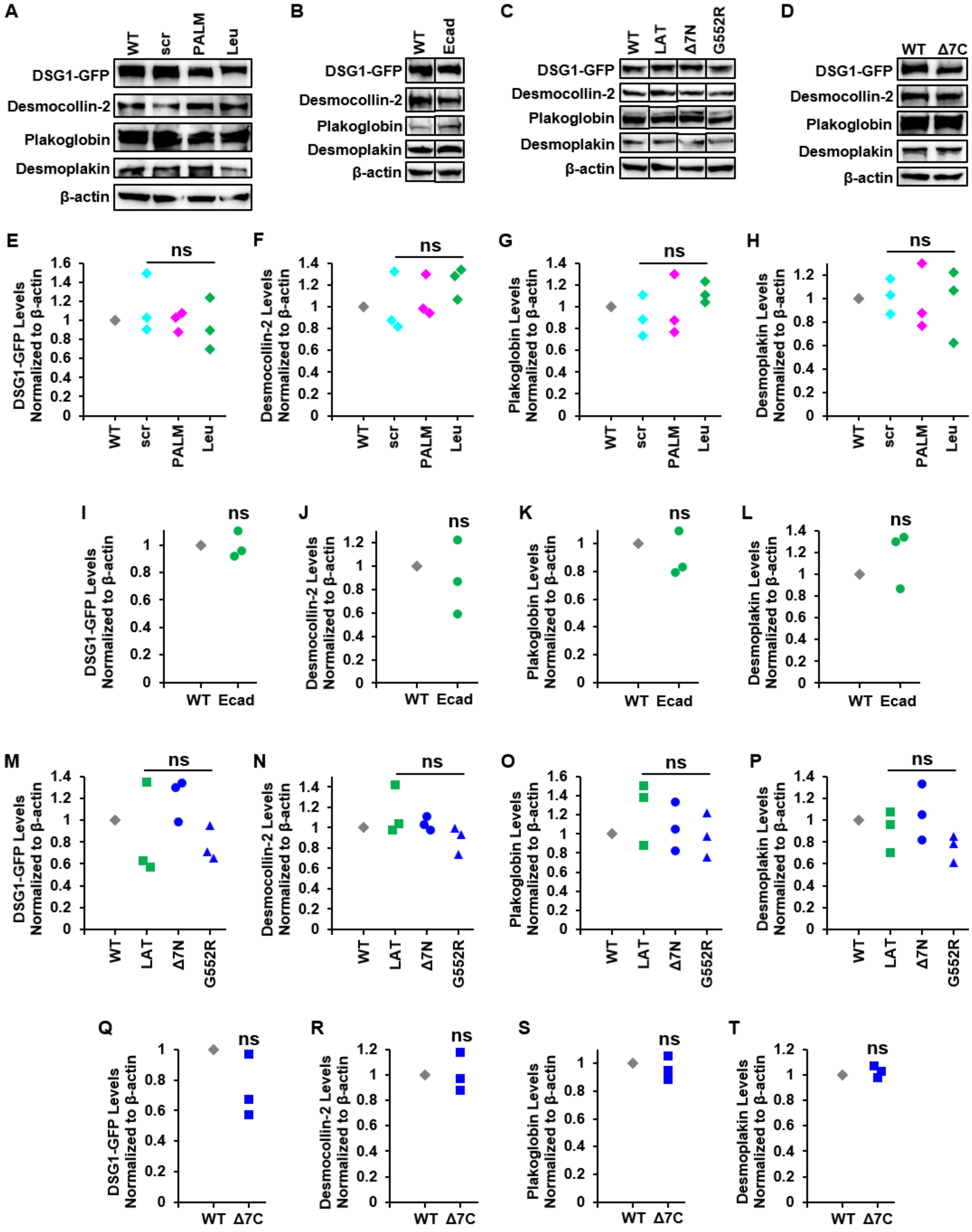

**Supplemental Figure 1:** DSG1-TMD-GFP variants are expressed at similar levels and do not impact expression levels of other desmosomal proteins. (A-D) Western blots show expression levels of DSG1-GFP, desmocollin-2, plakoglobin, or desmoplakin alongside β-actin in populations of DSG-null cells expressing $DSG1_{WT}$-GFP or $DSG1_{TMD}$-GFP variants. Cell lines were generated in four groups and expression levels were checked for each group prior to conducting further experiments. Group 1 includes $DSG1_{scr}$-GFP, $DSG1_{PALM}$-GFP, and $DSG1_{Leu}$-GFP. Group 2 includes $DSG1_{Ecad}$. $DSG1_{WT}$-GFP and $DSG1_{Ecad}$-GFP bands come from the same cropped gel which contained additional samples not shown in this work; crops are indicated by vertical lines. Group 3 includes $DSG1_{LAT}$-GFP, $DSG1_{\Delta 7N}$-GFP, and $DSG1_{G552R}$-GFP. These bands are from the same cropped gel run in duplicate alongside an additional cell line not reported in this work; crops are indicated by vertical lines. Group 4 includes $DSG1_{\Delta 7C}$-GFP. These bands were run alongside additional samples not shown in this work which were cropped out. $DSG1_{SAM}$-GFP is not shown as it was previously published (Zimmer et al 2021). (E-T) Quantification of bands in (A-D) showing DSG1-GFP, desmocollin-2, plakoglobin, and desmoplakin normalized to β-actin: Group 1 (E-H), Group 2 (I-L), Group 3 (M-P), and Group 4 (Q-T).

**Supplemental Figure 2**

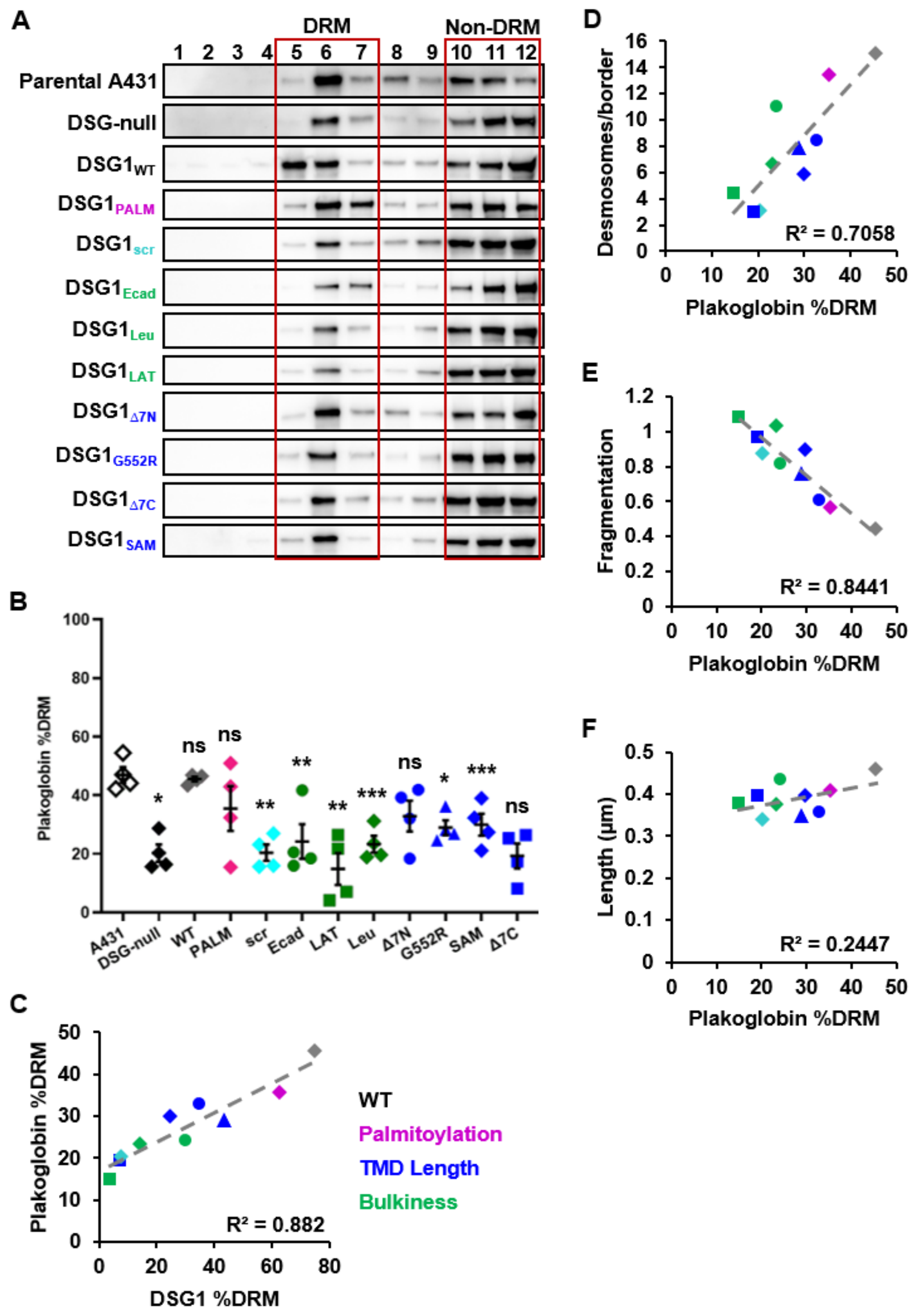


**Supplemental Figure 2:** Plakoglobin raft association depends on DSG1. (A) Sucrose gradient fractionations from DSG-null cells expressing $DSG1_{WT}$-GFP or each of the $DSG1_{TMD}$-GFP variants show distribution of plakoglobin between DRM and non-DRM fractions. (B) Quantification of westerns in (A). Error bars show mean +/- SEM, n = 4. (C) Scatter plot of correlation between DSG1 raft association and plakoglobin raft association, color-coded by TMD property. Point shapes and colors match those used in (B). (D-F) Scatter plots of correlations between PG raft association and desmosome number (D), desmosome length (E), and monolayer fragmentation (F).

**Supplemental Figure 3**

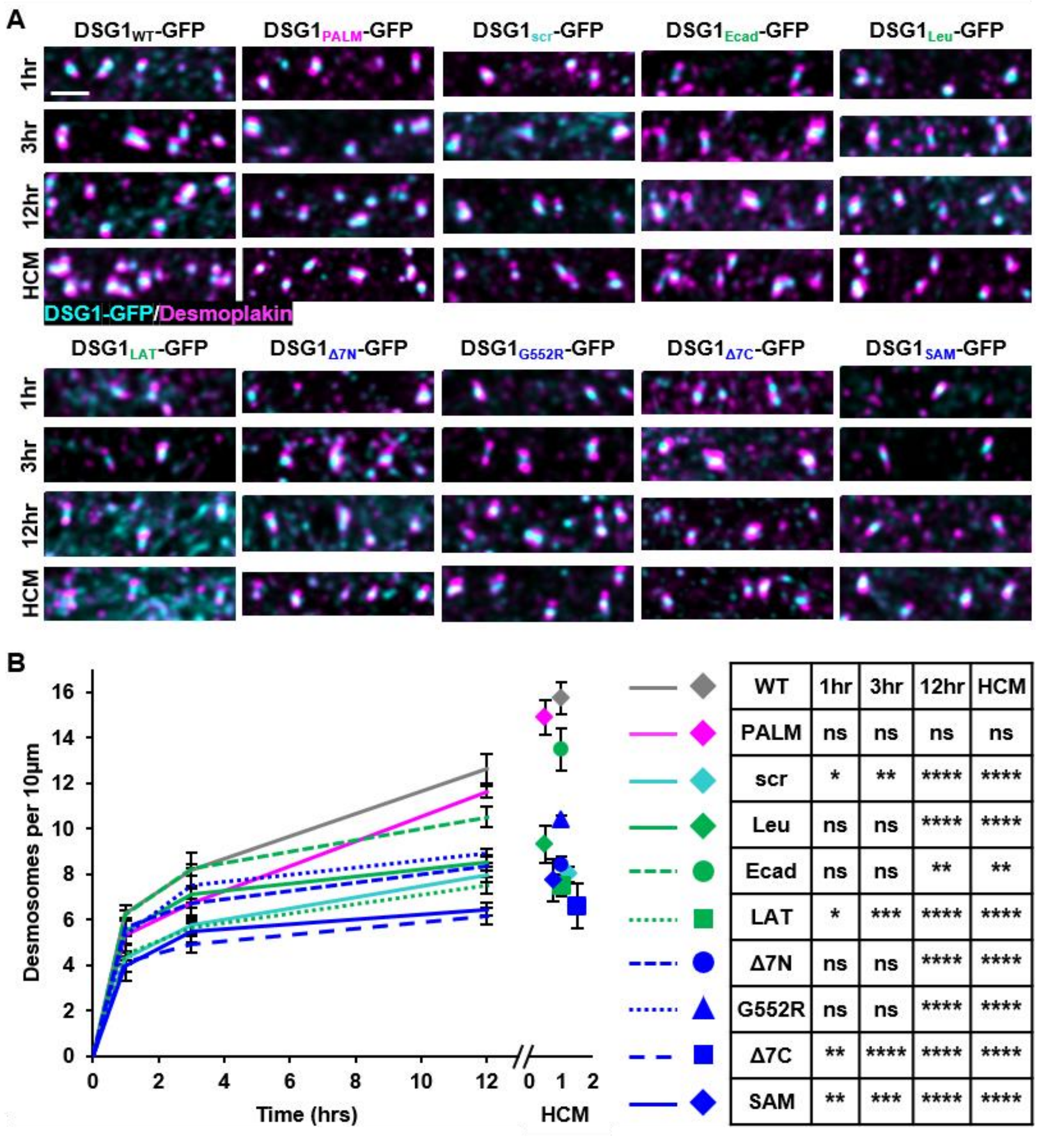


| WT | 1hr | 3hr | 12hr | HCM |
|---|---|---|---|---|
| PALM | ns | ns | ns | ns |
| scr | * | ** | **** | **** |
| Leu | ns | ns | **** | **** |
| Ecad | ns | ns | ** | ** |
| LAT | * | *** | **** | **** |
| Δ7N | ns | ns | **** | **** |
| G552R | ns | ns | **** | **** |
| Δ7C | ** | **** | **** | **** |
| SAM | ** | *** | **** | **** |

**Supplemental Figure 3:** DSG1 raft association regulates desmosome dynamics. (A) Spinning disk confocal images of DSG-null cells expressing DSG1$_{WT}$-GFP or other DSG1$_{TMD}$-GFP variants stained for GFP and DP at steady state or 1, 3, or 12 hours after a calcium switch. Bar, 1 μm. (B) Graph showing desmosome assembly progression, measured in desmosome quantity, following a calcium switch and compared to desmosome quantities at steady state.

**Supplemental Figure 4**

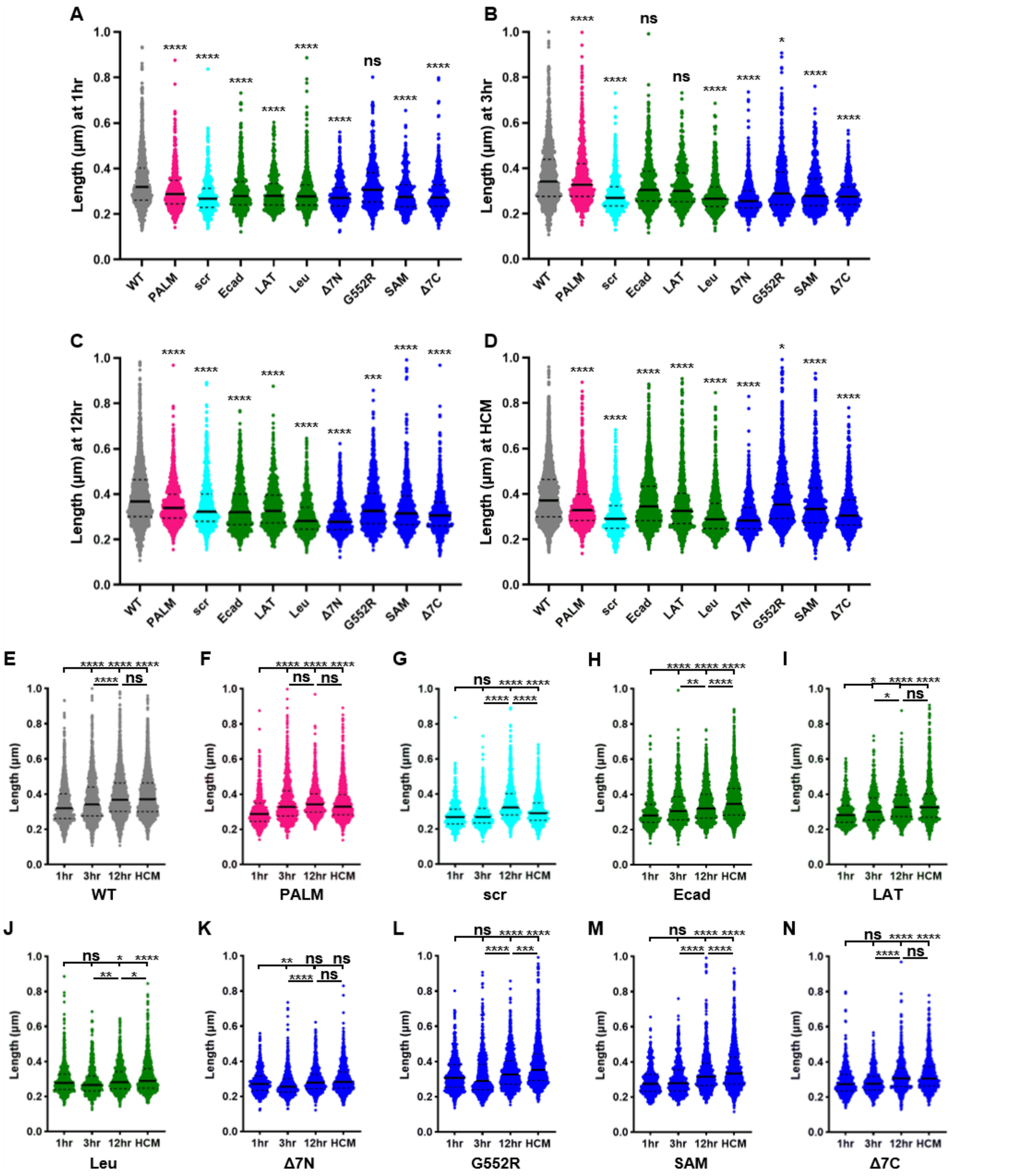


**Supplemental Figure 4:** Non-raft DSG1$_{TMD}$ variants form smaller desmosomes during assembly. (A-D) Violin plots show distribution of desmosome lengths after 1hr (A), 3hr (B), or 12hr (C) calcium switch or at steady state (HCM, D) for all DSG1$_{TMD}$-GFP variants. E-N) Violin plots show distribution of desmosome lengths for cells expressing DSG1$_{WT}$-GFP (E), DSG1$_{PALM}$-GFP (F), DSG1$_{scr}$-GFP (G), DSG1$_{Ecad}$-GFP (H), DSG1$_{LAT}$-GFP (I), DSG1$_{Leu}$-GFP (J), DSG1$_{\Delta 7N}$-GFP (K), DSG1$_{G552R}$-GFP (L), DSG1$_{SAM}$-GFP (M), DSG1$_{\Delta 7C}$-GFP (N). Desmosome length measurements come from 3 independent replicates where each replicate included 10 images per $Dsg1_{TMD}$ variant per timepoint. Each image contained the borders of at least one cell and its neighbors. All desmosomes found in an image were counted and measured amounting to 20-300+ desmosomes per image depending on timepoint and expressed variant. Y-axis was limited to 1 µm to better visualize and compare spreads. The following variants at conditions had limited measurements above 1 µm: $DSG1_{WT}$-GFP at 3hr, 12hr, and HCM, $DSG1_{PALM}$-GFP at 3hr and HCM, $DSG1_{Ecad}$-GFP at 12hr and HCM, $DSG1_{LAT}$-GFP at HCM, $DSG1_{G552R}$-GFP at 12hr and HCM, and $DSG1_{SAM}$-GFP at 12hr and HCM.

**Supplemental Figure 5**

A

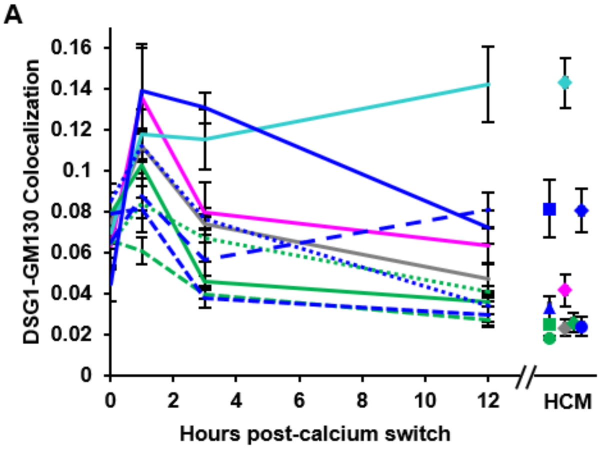


| WT | LCM | 1hr | 3hr | 12hr | HCM |
|---|---|---|---|---|---|
| PALM | ns | ns | ns | ns | ns |
| scr | ns | ns | * | **** | **** |
| Leu | ns | ns | ns | ns | ns |
| Ecad | ns | ns | ns | ns | ns |
| LAT | ns | ns | ns | ns | ns |
| Δ7N | ns | ns | ns | ns | ns |
| G552R | ns | ns | ns | ns | ns |
| Δ7C | ns | ns | ns | ns | ** |
| SAM | ns | ns | ** | ns | ** |

B

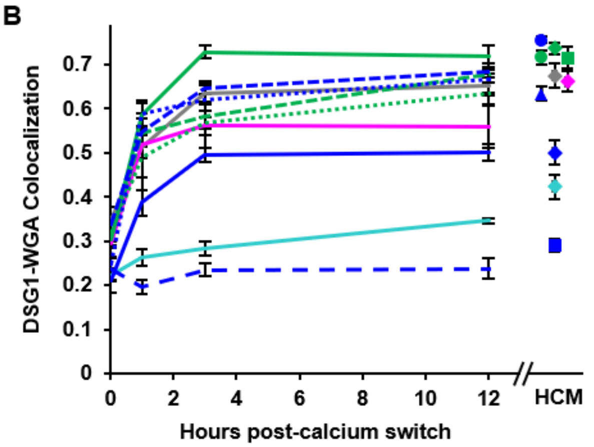


| WT | LCM | 1hr | 3hr | 12hr | HCM |
|---|---|---|---|---|---|
| PALM | ns | ns | ns | ns | ns |
| scr | * | **** | **** | **** | **** |
| Leu | ns | ns | ns | ns | ns |
| Ecad | ns | ns | ns | ns | ns |
| LAT | ns | ns | ns | ns | ns |
| Δ7N | ns | ns | ns | ns | ns |
| G552R | ns | ns | ns | ns | ns |
| Δ7C | ns | **** | **** | **** | **** |
| SAM | * | * | * | ** | ** |

**Supplemental Figure 5:** Most DSG1$_{TMD}$-GFP variants exhibit normal trafficking with substantial membrane-localized pools. (A) Colocalization between DSG1-GFP and GM130. (B) Colocalization between DSG1-GFP and WGA. Error bars represent mean +/- SEM, n = 3. (C-L) Images show DSG-null cells expressing DSG1$_{TMD}$-GFP variants stained for GFP, WGA, and GM130 at steady state or after a 1-, 3-, or 12-hour calcium switch: DSG1$_{WT}$-GFP (C), DSG1$_{PALM}$-GFP (D), DSG1$_{scr}$-GFP (E), DSG1$_{Ecad}$-GFP (F), DSG1$_{LAT}$-GFP (G), DSG1$_{Leu}$-GFP (H), DSG1$_{\Delta 7N}$-GFP (I), DSG1$_{G552R}$-GFP (J), DSG1$_{SAM}$-GFP (K), DSG1$_{\Delta 7C}$-GFP (L). Images on subsequent pages. Bar, 5µm.

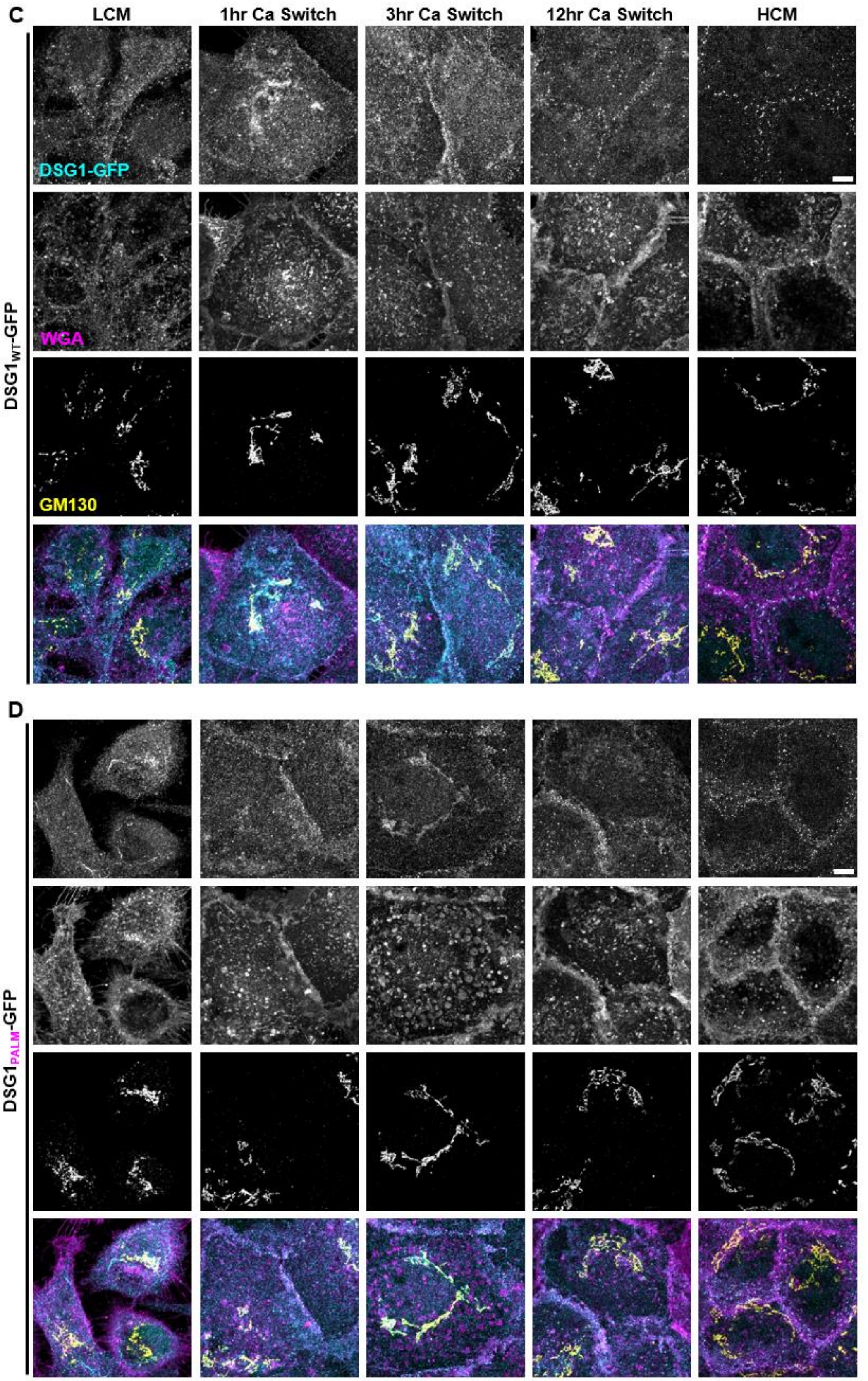
C
LCM
1hr Ca Switch
3hr Ca Switch
12hr Ca Switch
HCM
DSG1-GFP
WGA
GM130
DSG1$_{WT}$-GFP
D
DSG1$_{PALM}$-GFP

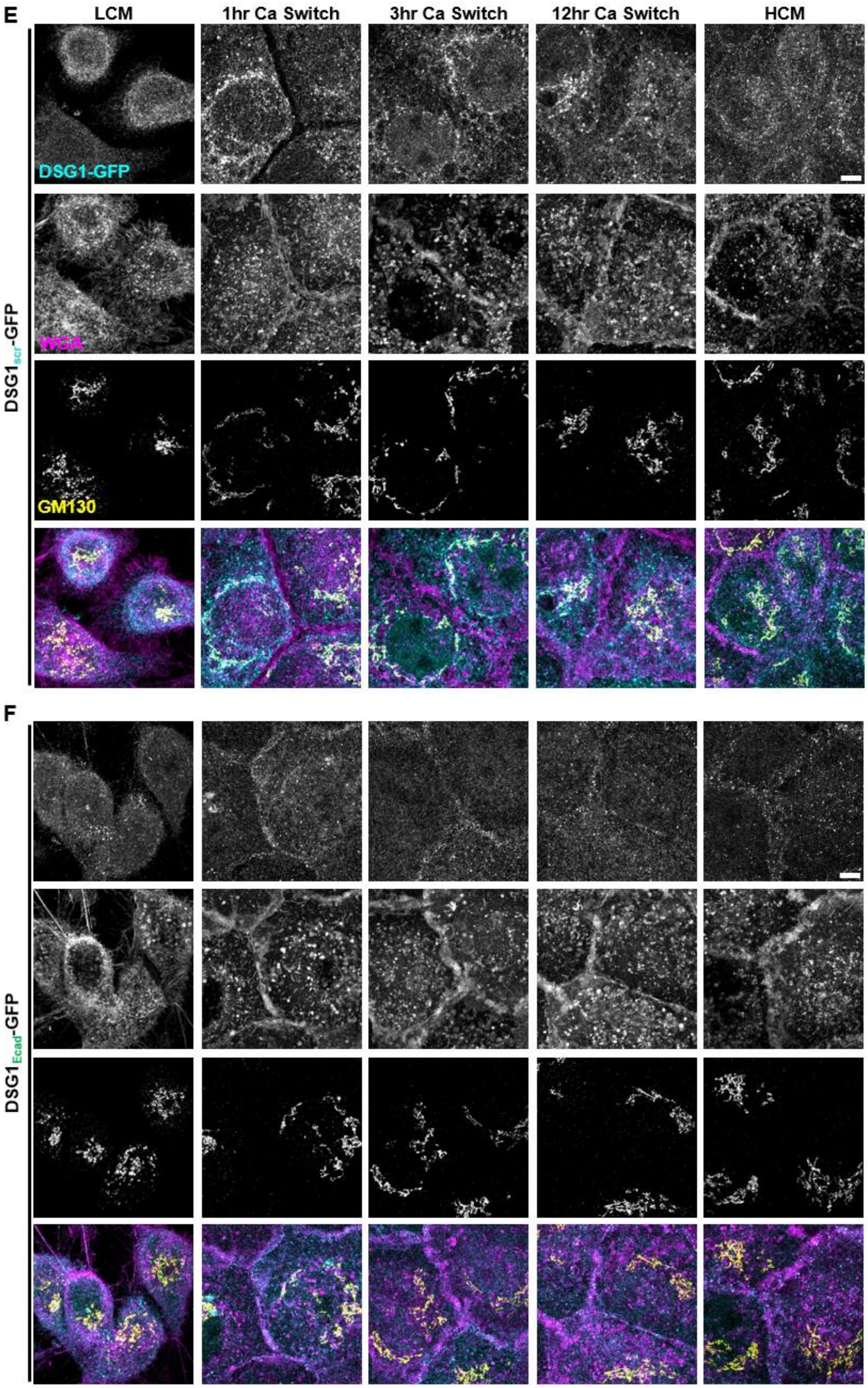
E
LCM
1hr Ca Switch
3hr Ca Switch
12hr Ca Switch
HCM
DSG1-GFP
WGA
GM130
DSG1scr-GFP
F
DSG1Ecad-GFP

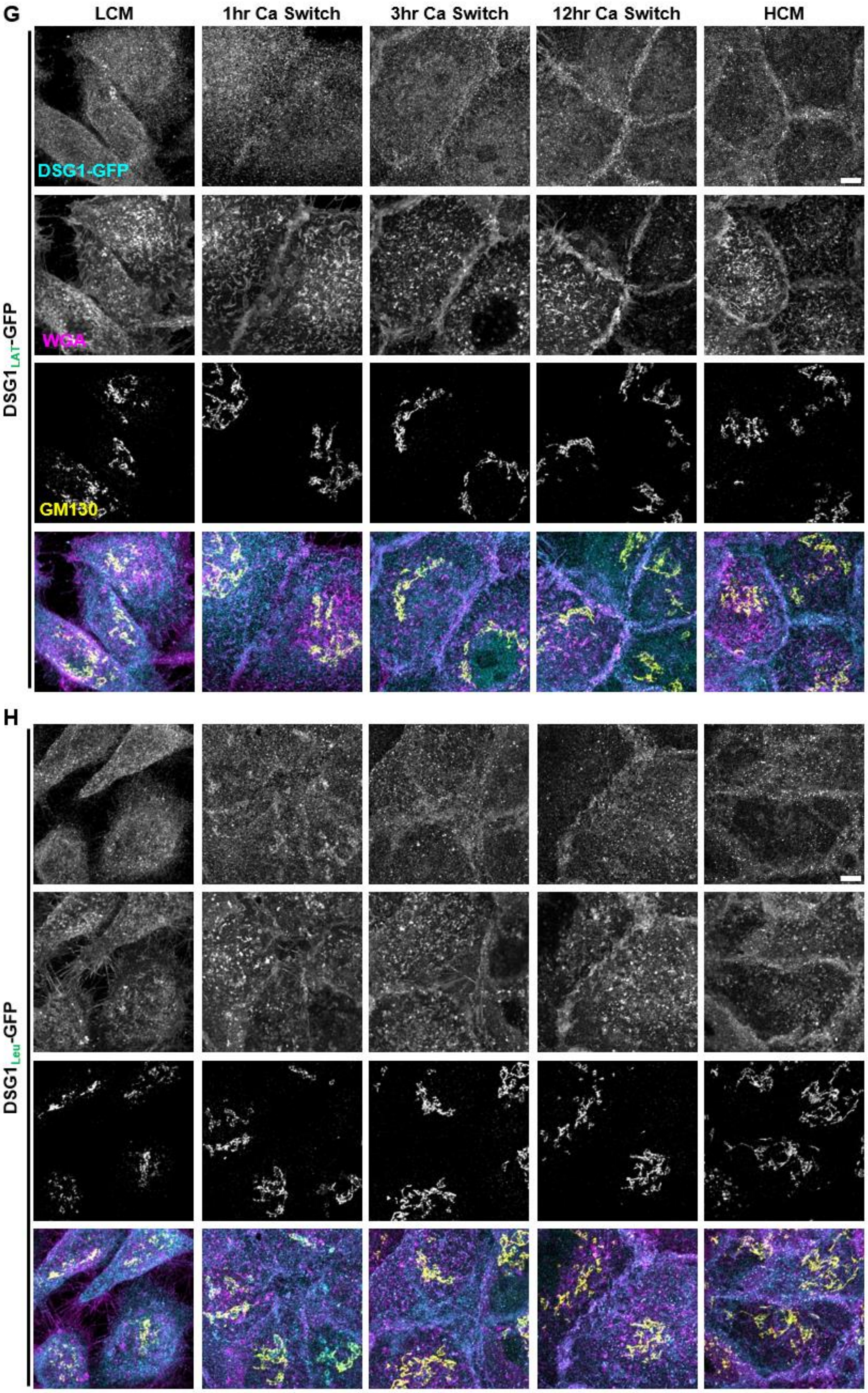

G
LCM
1hr Ca Switch
3hr Ca Switch
12hr Ca Switch
HCM
DSG1-GFP
WGA
GM130
DSG1LAT-GFP
H
DSG1Leu-GFP

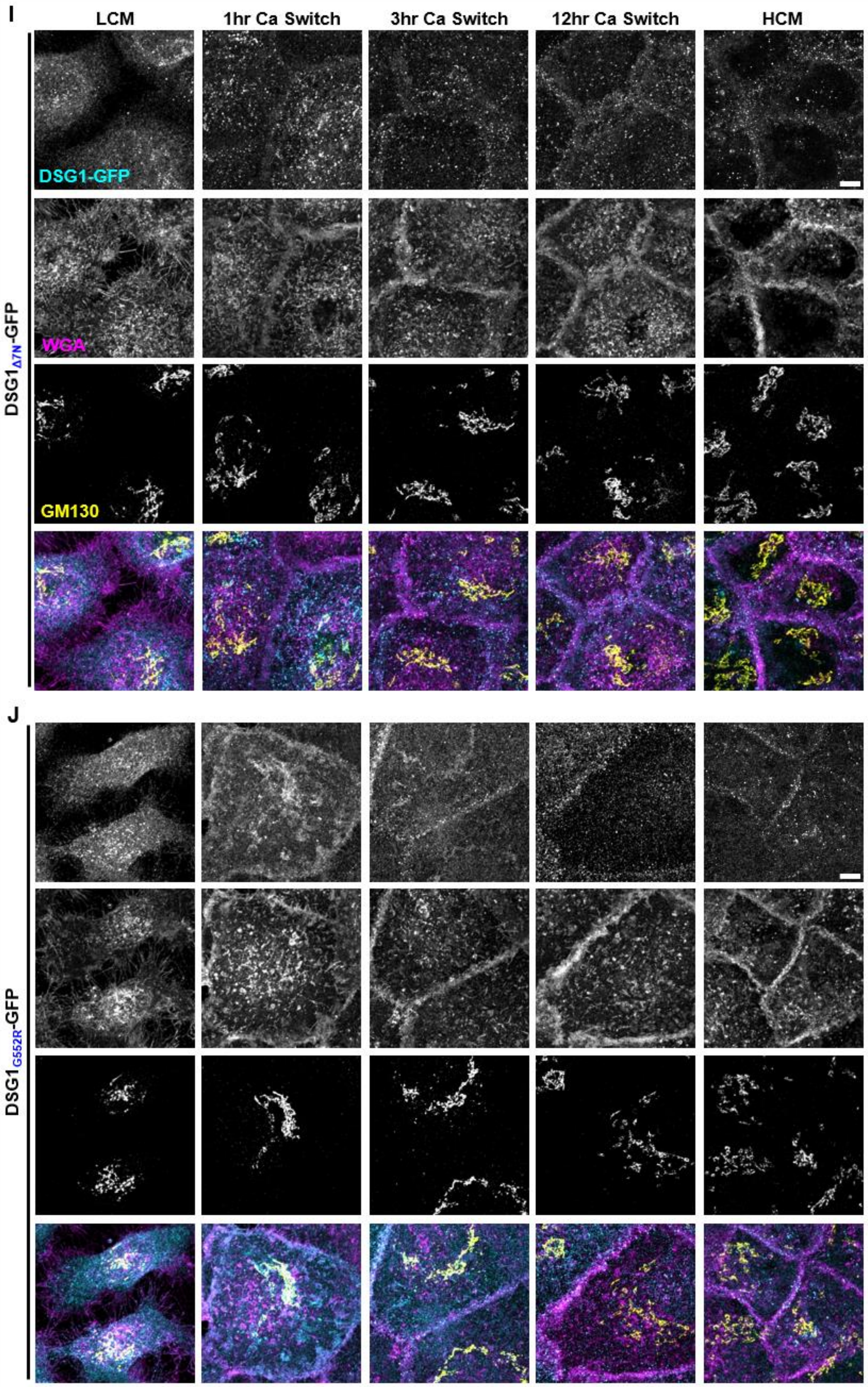
I
LCM
1hr Ca Switch
3hr Ca Switch
12hr Ca Switch
HCM
DSG1-GFP
WGA
GM130
DSG1Δ7N-GFP
J
DSG1G552R-GFP

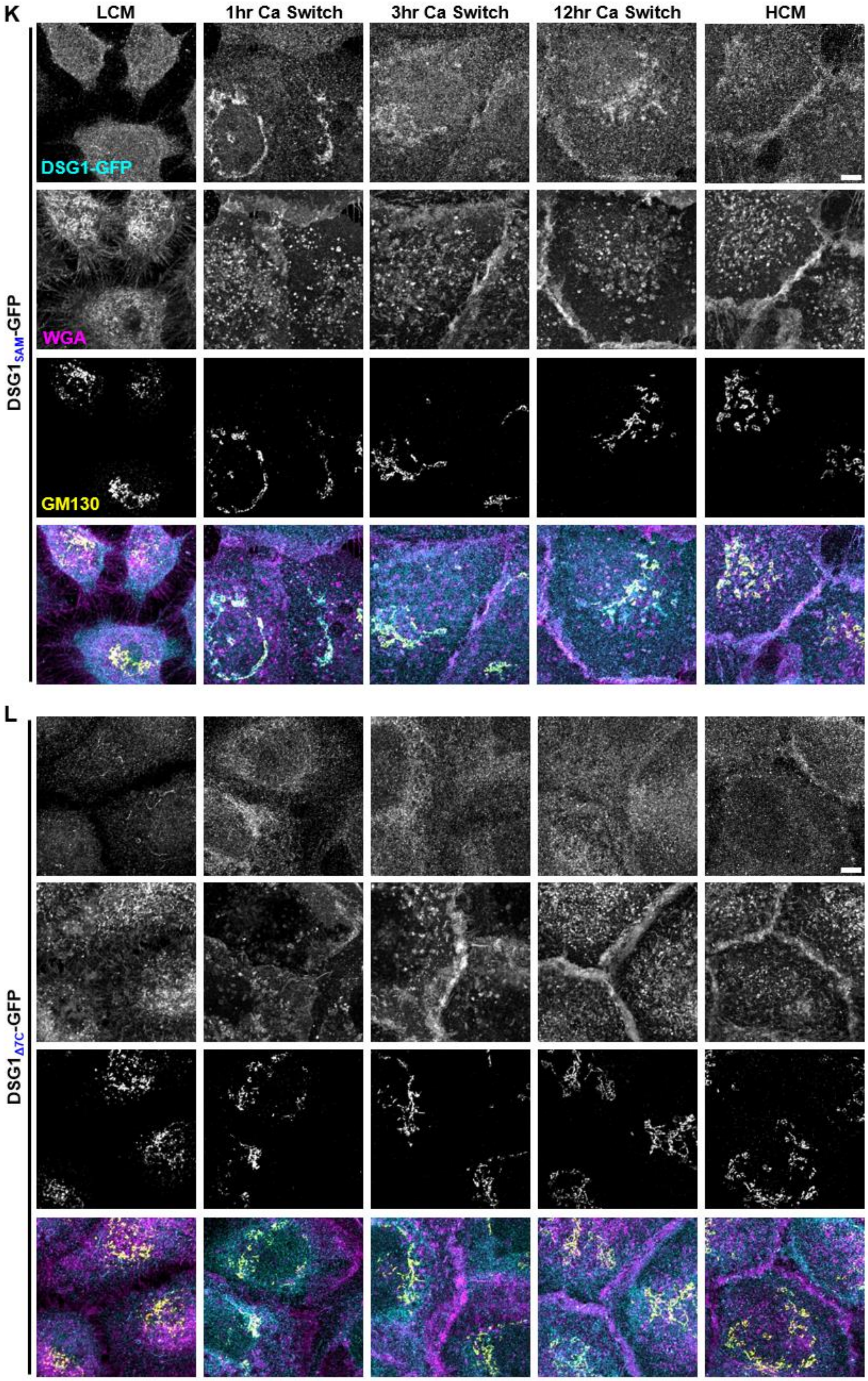

K
LCM
1hr Ca Switch
3hr Ca Switch
12hr Ca Switch
HCM
DSG1-GFP
WGA
GM130
DSG1SAM-GFP
L
DSG1Δ7C-GFP

**Supplemental Figure 6**

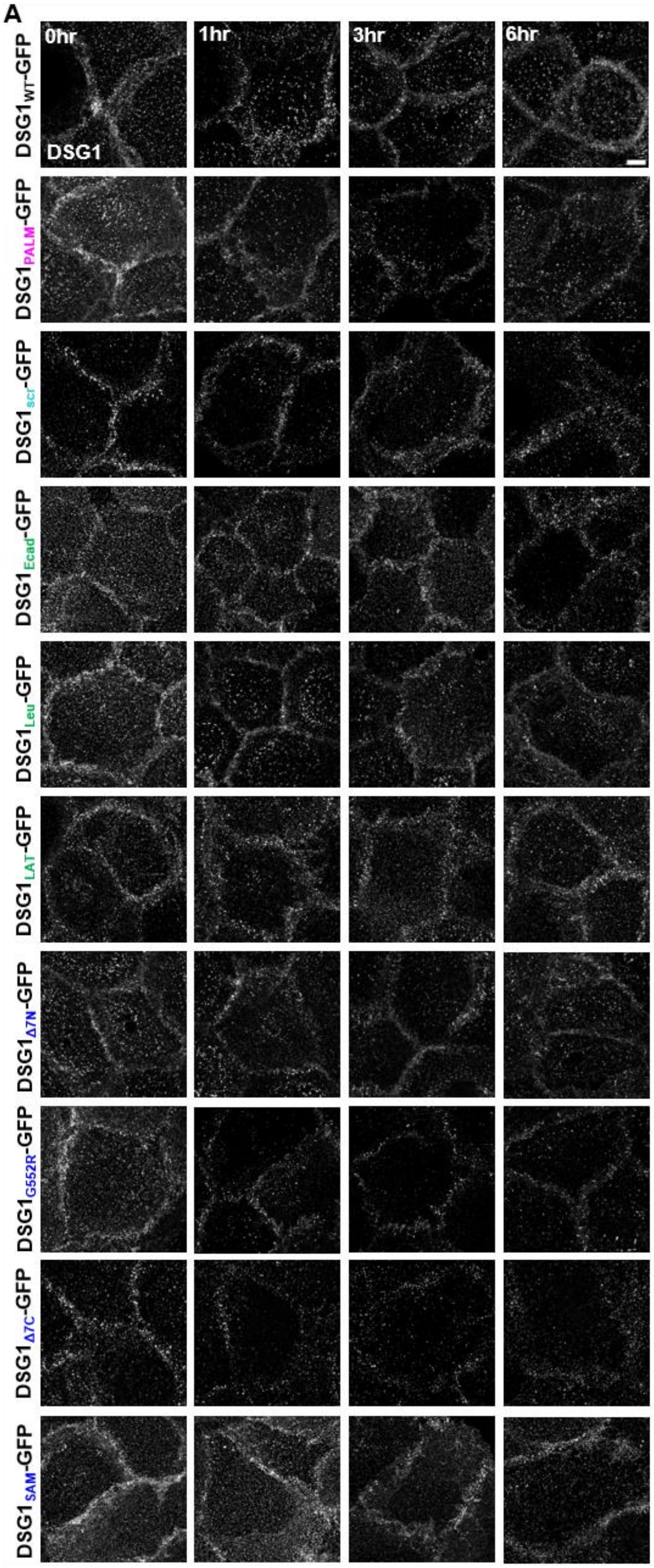


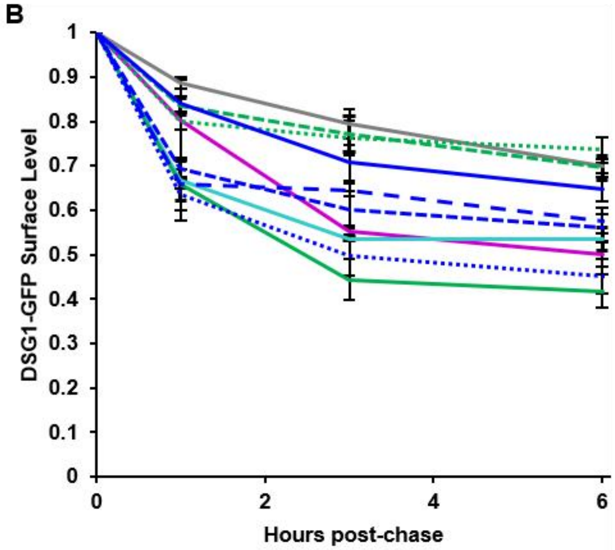


| WT | PALM | scr | Leu | Ecad | LAT | Δ7N | G552R | Δ7C | SAM |
|---|---|---|---|---|---|---|---|---|---|
| 1hr | ns | **** | **** | ns | ns | *** | **** | **** | ns |
| 3hr | **** | **** | **** | ns | ns | *** | **** | ** | ns |
| 6hr | *** | ** | **** | ns | ns | ** | **** | * | ns |

**Supplemental Figure 6**: Many $DSG1_{TMD}$-GFP variants exhibit increased surface turnover. (A) Images show surface level DSG1 in DSG-null cells expressing $DSG1_{TMD}$-GFP variants 0, 1, 3, or 6 hours after pulse-chase with antibody against DSG1 extracellular domain. Bar, 5 µm. (B) Quantification of images in (A). Error bars represent mean +/- SEM, n = 3.

**Supplemental Figure 7**

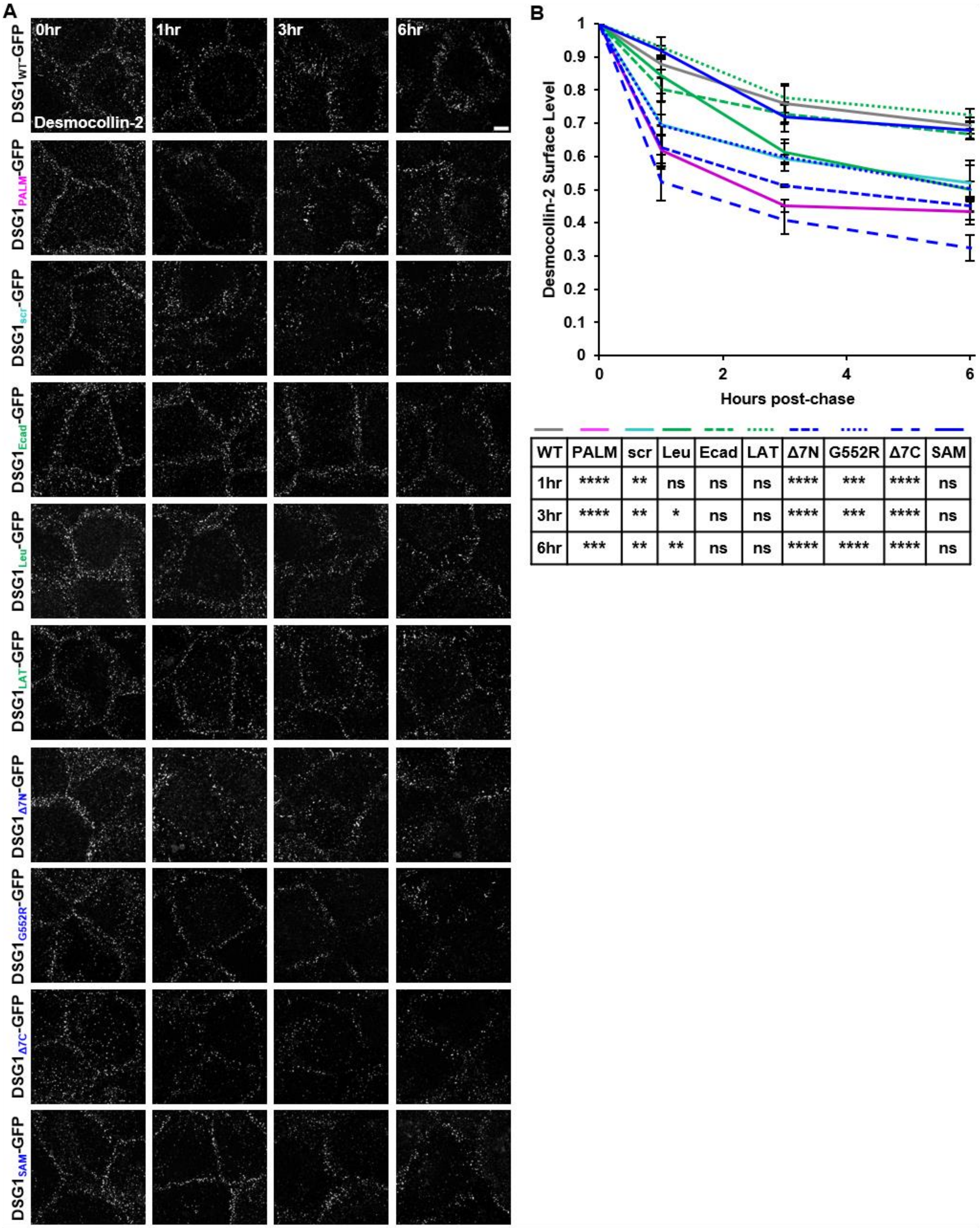


| WT | PALM | scr | Leu | Ecad | LAT | Δ7N | G552R | Δ7C | SAM |
|---|---|---|---|---|---|---|---|---|---|
| 1hr | **** | ** | ns | ns | ns | **** | *** | **** | ns |
| 3hr | **** | ** | * | ns | ns | **** | *** | **** | ns |
| 6hr | *** | ** | ** | ns | ns | **** | **** | **** | ns |

**Supplemental Figure 7:** DSC2 surface turnover mirrors that of Dsg1-GFP. (A) Images show endogenous surface level DSC2 in DSG-null cells expressing DSG1$_{TMD}$-GFP variants 0, 1, 3, or 6 hours after pulse-chase with antibody against extracellular DSC2 domain. Bar, 5 µm. (B) Quantification of images in (A). Error bars represent mean +/- SEM, n = 3.
